# Mechanism of Viral Glycoprotein Targeting by Membrane-associated-RING-CH Proteins

**DOI:** 10.1101/2021.01.25.428025

**Authors:** Cheng Man Lun, Abdul A. Waheed, Alhlam Majadly, Nicole Powell, Eric O. Freed

## Abstract

An emerging class of cellular inhibitory proteins has been identified that targets viral glycoproteins. These include the membrane-associated RING-CH (MARCH) family of E3 ubiquitin ligases that, among other functions, downregulate cell-surface proteins involved in adaptive immunity. The RING-CH domain of MARCH proteins is thought to function by catalyzing the ubiquitination of the cytoplasmic tails (CTs) of target proteins, leading to their degradation. MARCH proteins have recently been reported to target retroviral envelope glycoproteins (Env) and vesicular stomatitis virus G glycoprotein (VSV-G). However, the mechanism of antiviral activity remains poorly defined. Here we show that MARCH8 antagonizes the full-length forms of HIV-1 Env, VSV-G, Ebola virus glycoprotein (EboV-GP), and the spike (S) protein of severe acute respiratory syndrome coronavirus-2 (SARS-CoV-2) thereby impairing the infectivity of virions pseudotyped with these viral glycoproteins. This MARCH8-mediated targeting of viral glycoproteins requires the E3 ubiquitin ligase activity of the RING-CH domain. We observe that MARCH8 protein antagonism of VSV-G is CT dependent. In contrast, MARCH8-mediated targeting of HIV-1 Env, EboV-GP and SARS-CoV-2 S protein by MARCH8 does not require the CT, suggesting a novel mechanism of MARCH-mediated antagonism of these viral glycoproteins. Confocal microscopy data demonstrate that MARCH8 traps the viral glycoproteins in an intracellular compartment. We observe that the endogenous expression of *MARCH8* in several relevant human cell types is rapidly inducible by type I interferon. These results help to inform the mechanism by which MARCH proteins exert their antiviral activity and provide insights into the role of cellular inhibitory factors in antagonizing the biogenesis, trafficking, and virion incorporation of viral glycoproteins.

**Importance:** Viral envelope glycoproteins are an important structural component on the surface of enveloped viruses that direct virus binding and entry and also serve as targets for the host adaptive immune response. In this study, we investigate the mechanism of action of the MARCH family of cellular proteins that disrupt the trafficking and virion incorporation of viral glycoproteins across several virus families. This research provides novel insights into how host cell factors antagonize viral replication, perhaps opening new avenues for therapeutic intervention in the replication of a diverse group of highly pathogenic enveloped viruses.

## INTRODUCTION

Viruses rely heavily on host cellular machinery to replicate due to their relatively limited coding capacity. In turn, cells have evolved extensive and elaborate defense mechanisms to impede virus replication. Cells encode numerous proteins, often referred to as inhibitory or restriction factors, that are central components of the innate immune response that serves as the first line of defense against invading pathogens. Well-characterized restriction factors include apolipoprotein B mRNA editing catalytic polypeptide-like (APOBEC) family proteins, tripartite motif protein 5 α, (TRIM5α), SAM and HD domain-containing protein 1 (SAMHD1), Myxovirus resistance 2 (Mx2), and tetherin/BST-2 (1). These restriction factors are either expressed constitutively and/or are induced by IFN (2–4). Recently, an emerging class of host proteins have been shown to specifically target the synthesis, trafficking and/or function(s) of viral glycoproteins (5). These include the IFN-induced transmembrane (IFITM) proteins, guanylate-binding proteins (GBPs), endoplasmic reticulum class I α-mannosidase (ERManI), galectin 3 binding protein (LGALS3BP/90K), Ser incorporator (SERINC), and the MARCH proteins (5–13).

Viral envelope glycoproteins, which are important structural components decorating the surface of enveloped virus particles, recognize receptors on target cells and mediate viral entry by catalyzing membrane fusion events either at the plasma membrane or in low-pH endosomes following endocytic uptake of the viral particle. They are synthesized and cotranslationally glycosylated in the ER and then traffic through the secretory pathway to the site of virus assembly (14). During trafficking through the Golgi apparatus, some viral glycoproteins are cleaved by furin or furin-like proteases as a requisite step in the generation of the fusion-active viral glycoprotein complex. Viral glycoproteins often multimerize (e.g., as trimers) during trafficking (14). Many enveloped viruses assemble at the plasma membrane, whereas some assemble in alternative compartments. For example, SARS-CoV-2 particles are thought to assemble in the ER/Golgi intermediate compartment (ERGIC) and be released by exploiting lysosomal organelles (15, 16).

Ubiquitin is a small, 76-amino acid protein that is highly expressed in eukaryotic cells. It can be attached to target proteins, usually on Lys residues, but occasionally on Ser, Thr, or Cys residues, via a multi-enzyme cascade involving an E1 ubiquitin-activating enzyme, an E2 ubiquitin-conjugating enzyme, and an E3 ubiquitin ligase (17). Ubiquitin attachment can serve as a signal for target protein degradation in the proteasome or lysosome, or can regulate the endocytic trafficking of the target protein or other aspects of protein function.

The MARCH family of RING-finger E3 ubiquitin ligases comprise 11 structurally diverse members. With the exception of the cytosolic MARCH7 and MARCH10 proteins, MARCH family members are transmembrane proteins containing multiple (ranging from 2-14) putative membrane-spanning domains and bearing an N-terminal cytoplasmic RING-CH domain (Fig. 1) (18). MARCH proteins were originally discovered as cellular homologs of the K3 and K5 E3 ubiquitin ligases of Kaposi’s sarcoma-associated herpes virus (KSHV) (19, 20). These viral MARCH protein homologs confer escape from the host immune response by downregulating the major histocompatibility class I (MHC-I) antigen on the surface of virus-infected cells (21, 22). Other large DNA viruses encode analogous proteins involved in immune evasion (23). Cellular MARCH proteins have been reported to downregulate numerous proteins, including MHC-II (24), transferrin receptor (25), TRAIL receptors (26), CD44 and CD81 (27), IL-1 receptor accessory protein (28), CD98 (29), and tetherin/BST-2 (30). The RING-CH domain of MARCH proteins and its interaction with E2 enzymes are responsible for their E3 ligase activity; thus, mutations that either inactivate the catalytic center of the RING domain, or prevent interaction with E2s, abrogate MARCH protein activity (18, 25, 28). MARCH proteins and their viral homologs generally catalyze the transfer of ubiquitin to Lys residues in the CTs of their target proteins, leading to their degradation and/or altered trafficking (18, 19, 25, 31–34). However, the viral MARCH protein homologs K3 and K5 of KSHV have been reported to attach ubiquitin to Ser, Thr, and/or Cys residues (35–37).

**Figure 1.**
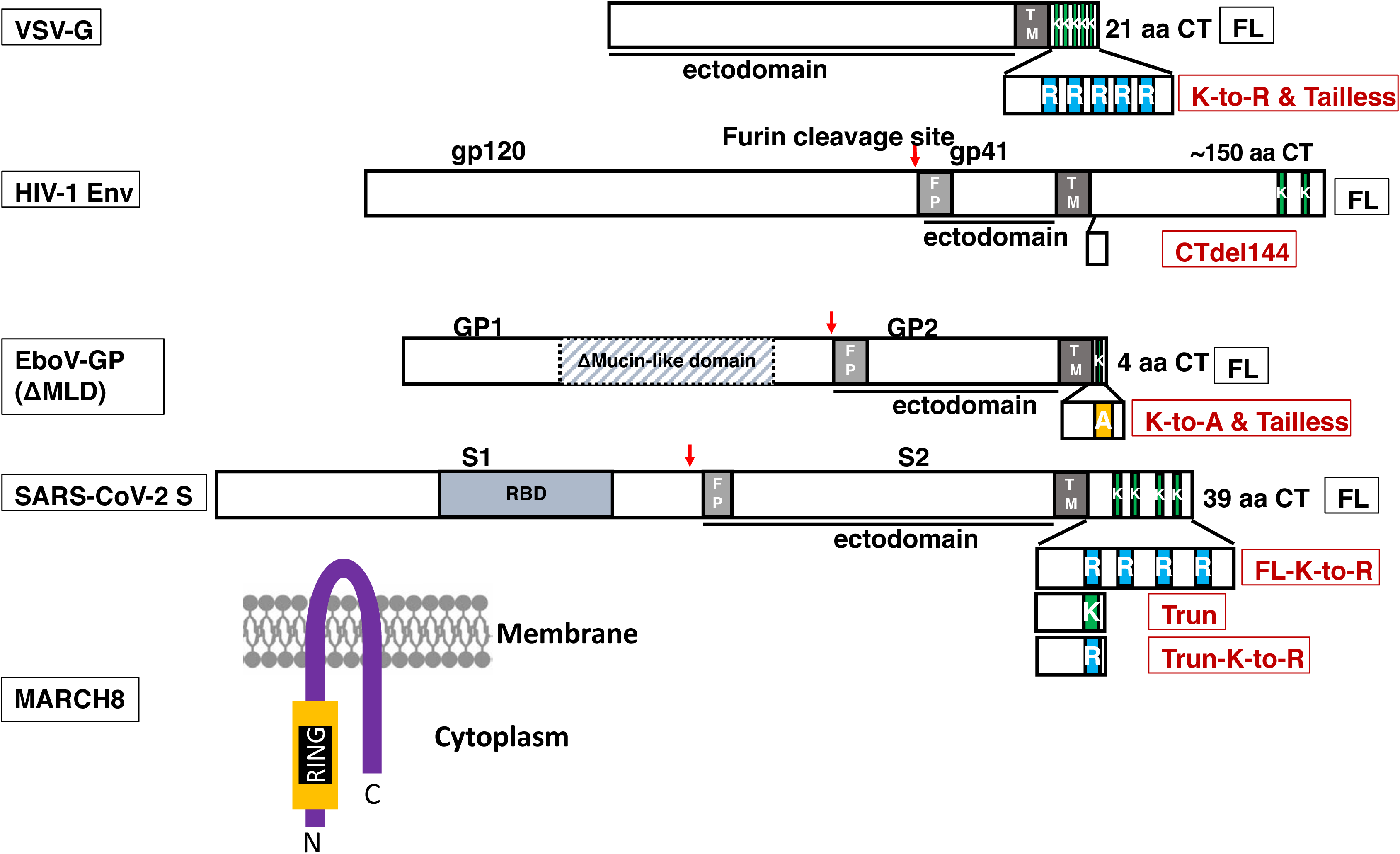
Organization of the viral envelope glycoproteins used in this study. Surface and transmembrane subunits of HIV-1 Env (gp120 and gp41), EboV-GP (GP1 and GP2) and SARS-CoV-2 S protein (S1 and S2) are indicated. Where appropriate, the site of furin cleavage is indicated by the red arrow. The Δmucin-like domain (ΔMLD, dotted rectangle with hatching mark) and the receptor-binding domain (RBD) of the EboV-GP and SARS-CoV-2 S protein, respectively, are indicated. The fusion peptide (FP) and transmembrane domains (TM) are labeled. The length in amino acids (aa) of each full-length (FL) CT, and the number of Lys (K) residues in each CT, are indicated. Each CT mutant, and the Lys residues mutated to either Arg (R) or Ala (A), are shown. At the bottom is shown a schematic representation of the MARCH8 topology in the membrane, with two transmembrane domains and the RING finger E3 ligase domain. The structures of the viral glycoproteins and MARCH8 are not drawn to scale.

MARCH8 was first identified in a genome-wide siRNA screen as a potential HIV-1 restriction factor acting early in the virus replication cycle by an unknown mechanism (38). More recently, three MARCH proteins – MARCH1, 2 and 8 – were reported to be late-acting restriction factors that target retroviral Env glycoproteins and VSV-G, thereby impairing infectivity of HIV-1 virions bearing these viral glycoproteins (6, 8, 11). The endogenous expression of MARCH proteins differs among cell types, with higher expression of MARCH1, 2 and 8 observed in myeloid cells such as monocyte-derived macrophages (MDM) and monocyte-derived dendritic cells (MDDCs) compared to primary CD4^+^ T cells (6, 11). MARCH8 expression has been reported to be particularly high in the lung (19). MARCH8 knockdown or knockout in myeloid cells increases HIV-1 infectivity, suggesting that MARCH8 may serve as an antiviral factor in these cell types (11). MARCH1, 2, and 8 are reported to be localized to lysosomes, endosomes and the plasma membrane (18, 19, 25).

To understand the mechanism of MARCH-mediated antiviral activity in greater detail, envelope glycoproteins from four families of viruses were selected for study: the retrovirus HIV-1, the rhabdovirus VSV, the filovirus Ebola virus (EboV) and the coronavirus severe acute respiratory syndrome coronavirus-2 (SARS-CoV-2). Three of the glycoproteins encoded by these viruses [HIV-1 Env, EboV-GP, and SARS-CoV-2 spike (S) protein] are cleaved by furin, whereas VSV-G is not (39–41) (Fig.1). HIV-1 Env is synthesized as a precursor, gp160, that is cleaved to gp120 and gp41; the EboV-GP precursor pre-GP is cleaved to GP1 and GP2, and the SARS-CoV-2 S protein precursor is cleaved to S1 and S2 [Fig.1; (39–41)]. The CTs of the viral glycoproteins investigated in this study are highly variable in length, with those of HIV-1 Env, SARS-CoV-2 S protein, VSV-G, and EboV-GP containing 150, 39, 21 and 4 amino acids, respectively (**Fig. 1**). These viral glycoproteins have CTs that vary not only in length but also in the number of Lys residues they contain, with two Lys residues in the CT of HIV-1 Env, four in the SARS-CoV-2 S protein, five in VSV-G, and one in the CT of EboV-GP (Fig. 1). These Lys residues could potentially serve as targets for MARCH-mediated ubiquitination. Our data demonstrate that each of the viral glycoproteins examined is antagonized, to variable extents, by MARCH8. We observed that MARCH-mediated inhibition of VSV-G is CT dependent, whereas inhibition of HIV-1 Env, EboV-GP and SARS-CoV-2 S protein is CT independent. We further demonstrate that knock-down of endogenous *MARCH8* gene expression in HEK293T cells increases the infectivty of HIV-1 particles produced from those cells and that endogenous expression of *MARCH8* is induced by IFN treatment in a human T-cell line, hPBMCs, and primary human airway epithelial cells. Finally, we show that MARCH proteins colocalize with, and retain, the viral glycoproteins in an aberrant intracellular compartment that bears the lysosomal marker LAMP-1. Collectively, our data provide novel insights into the mechanism of action of the MARCH family of cellular E3 ubiquitin ligases and their ability to antagonize diverse viral envelope glycoproteins.

## RESULTS

### MARCH-mediated inhibition of viral envelope glycoproteins exhibits differential CT dependence

It has been shown that the ectopic expression of MARCH8 in virus-producer cells markedly reduces the infectivity of HIV-1 virions bearing retroviral Env glycoproteins or VSV-G (6, 11). However, the molecular mechanism by which MARCH8 targets viral glycoproteins is not well defined. As mentioned in the Introduction, it has been determined that MARCH proteins downregulate a number of proteins by transferring ubiquitin to their CTs, leading to their lysosomal degradation. To investigate the potential role of MARCH-mediated CT ubiquitination in the downregulation of viral glycoproteins, we deleted the CTs of HIV-1 Env, VSV-G, EboV-GP, and SARS-CoV-2 S protein (**Fig. 1**) and cotransfected the viral glycoprotein expression vectors with the Env(-) pNL4-3 derivative pNL4-3/KFS (42) or a luciferase-encoding NL4-3-derived vector virus. In the case of HIV-1, we used full-length pNL4-3 expressing WT Env or the CT-truncated Env mutant, CTdel-144 (43). Virus-containing supernatants were harvested, normalized for p24 capsid content or reverse transcriptase (RT) activity, and used to infect the TZM-bl indicator cell line (44) or, in the case of the S protein pseudotypes, HEK293T cells stably expressing the human angiotensin converting enzyme 2 (hACE2) receptor. Consistent with previous reports (6, 11), we observed that the infectivity of HIV-1 virions bearing VSV-G was markedly reduced (by ∼10-fold) upon expression of WT MARCH8 in the virus-producer cells (**Fig. 2A**). In contrast, the MARCH8-CS (25) and MARCH8-W114A (28, 45, 46) mutants, which abolish RING-CH function or interaction with E2 ubiquitin conjugating enzyme, respectively, did not exert antiviral activity (**Fig. 2A**). We observed that the infectivity conferred by VSV-G Tailless was ∼40% that of WT VSV-G, and overexpression of MARCH8 in virus-producer cells had no significant effect on the infectivity of this VSV-G variant (**Fig. 2A**). Next, we investigated the ability of MARCH8 to inhibit a VSV-G mutant in which the Lys residues in the CT, which are potential sites of MARCH8-mediated ubiquitination, were substituted. A VSV-G mutant in which the five Lys residues in the CT were mutated to Ala displayed severely compromised infectivity (data not shown). We therefore introduced the more conservative Lys-to-Arg substitution (K-to-R) at these five Lys residues (Fig. 1) and observed that this mutant is fully infectious relative to the WT. As observed with the VSV-G Tailless mutant, overexpression of MARCH8 in virus-producer cells had no significant effect on the infectivity of the VSV-G K-to-R mutant. As expected, the MARCH8 mutants likewise did not diminish the infectivity of either the Tailless or K-to-R VSV-G mutants. These results indicate that the ability of MARCH8 to antagonize VSV-G in this experimental system is dependent upon the Lys residues in the VSV-G CT, likely because they are targets of MARCH8-mediated ubiquitination.

**Figure 2.**
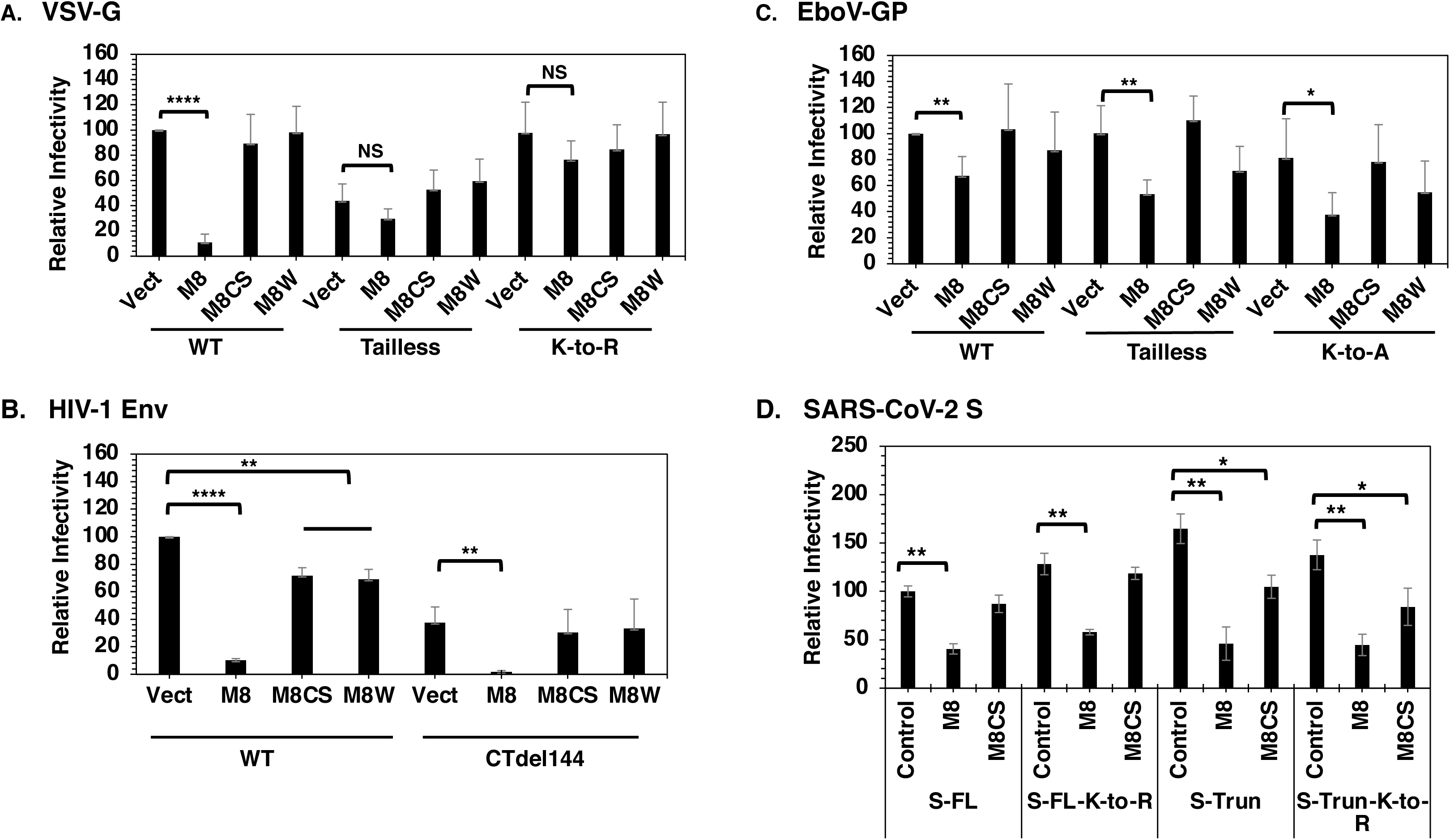
MARCH8-mediated targeting of viral envelope glycoproteins exhibits differential dependence on the CT. **(A)** HEK293T cells in a 12-well plate were cotransfected with the Env-defective HIV-1 molecular clone pNL4-3/KFS (1 μg) and vectors expressing WT or mutant VSV-G (100 ng) and HA-tagged WT or mutant MARCH8 (50 ng). TZM-bl cells were infected with VSV-G pseudotyped HIV-1 and luciferase activity was measured two days post-infection and normalized for viral p24. Infectivity of particles pseudotyped with WT VSV-G in the absence of MARCH8 is set to 100%. **(B)** HEK293T cells were cotransfected with WT or CT-deleted (CTdel144) pNL4-3 HIV-1 molecular clones (1 μg), and vectors expressing HA-tagged MARCH8 (WT or mutants) (100 ng). Two days post-transfection, virus supernatants were collected, analyzed for p24 content by western blot, and infections were carried out in TZM-bl cells, and normalized to p24 as described in **(A)**. Infectivity of WT HIV-1 in the absence of MARCH8 was set to 100%. **(C)** HEK293T cells were cotransfected with Env-defective (pNL4-3/KFS) HIV-1 molecular clone (1 μg), and vectors expressing EboV-GP (WT or CT mutants) (500 ng) and HA-tagged MARCH8 (WT or mutants) (250 ng), and infections were carried out in TZM-bl cells. Infectivity of particles pseudotyped with WT EboV-GP in the absence of MARCH8 was set to 100%. **D)** HEK293T cells in a 6-well plate were cotransfected with the Env-defective, luciferase-expressing pNL4-3 derivative pNL4-3.Luc.R-E- (3 μg) and vectors expressing SARS-CoV-2-S full-length (FL) and mutants (FL-K-to-R, Truncated, Truncated-K-to-R) (300 ng) and HA-tagged MARCH8 (WT or CS mutant) (200 ng). Two days post-transfection, virus supernatant was collected, and infectivity was measured in HEK293T cells stably expressing hACE2 with RT-normalized virus. Infectivity of virus particles pseudotyped with FL S protein in the absence of MARCH8 was set to 100%. Data shown are ± SD from 3-5 independent experiments. P values (two-tailed unpaired t-test): *p < 0.05, **p < 0.01, ****p < 0.0001.

As reported previously (6, 11), we observed that the infectivity of virus particles bearing HIV-1 Env is severely reduced (by ∼10-fold) upon overexpression of WT MARCH8 in the virus-producer cell. This inhibitory activity was to a large extent eliminated by the MARCH8-CS and MARCH8-W114A mutations (**Fig. 2B**). To investigate whether the inhibition of HIV-1 Env is CT dependent we tested the CTdel144 Env mutant (43), which lacks nearly the entire gp41 CT (**Fig. 1**). Under these assay conditions, the infectivity of CTdel144 HIV-1 was ∼40% that of WT HIV-1. Interestingly, in contrast to the lack of MARCH8-mediated inhibition observed with Tailless VSV-G, the infectivity of CT-deleted HIV-1 was reduced to a similar extent as the WT by expression of MARCH8 in the virus-producer cells. The MARCH8 mutants did not significantly inhibit the infectivity of the CTdel144 HIV-1 Env mutant. These results demonstrate that MARCH8-mediated antagonism of HIV-1 Env requires the E3 ubiquitin ligase activity of the RING domain but does not require the gp41 CT.

As indicated above, the dependency on the CT for MARCH8-mediated inhibition differs between VSV-G and HIV-1 Env. We next examined the effect of MARCH8 on the infectivity of the EboV-GP, which contains a CT of only four amino acids (**Fig. 1**). For these experiments, we used a version of the EboV-GP lacking the mucin-like domain (47) as we obtained higher infectivity with this variant (data not shown). We will refer to this delta mucin-like domain variant (ΔMLD) as WT because it contains an intact CT. The infectivity of HIV-1 virions bearing either the WT or CT-deleted EboV-GP was reduced by ∼30-40 % when the virus was produced in the presence of MARCH8. We also mutated the single Lys residue in the CT of EboV-GP and observed that the infectivity of this K-to-A mutant was inhibited by ∼50% in the presence of MARCH8. The infectivity of EboV-GP mutants was similar to that of the WT in the absence of MARCH8 overexpression (**Fig. 2C**). No statistically significant reduction in infectivity was observed for WT EboV-GP upon expression of the MARCH8-CS mutant, yet a minor reduction in infectivity was observed for EboV-GP Tailless and K-To-A mutants in the presence of the MARCH8-W114A mutant (**Fig. 2C**). These results demonstrate that MARCH8-mediated antagonism of EboV-GP is dependent on the E3 ubiquitin ligase activity of the RING domain but does not require the short EboV-GP CT.

We next tested the effect of MARCH8 overexpression on the infectivity of particles bearing the SARS-CoV-2 S protein, which has a 39 amino acid CT with five Lys residues. We also tested a CT-truncated SARS-CoV-2 S protein, which bears a 19-amino-acid truncation and contains one Lys residue (**Fig. 1**). The infectivity of HIV-1 virions bearing WT or CT-truncated SARS-CoV-2 S protein was reduced by ∼60% in the presence of MARCH8. We also mutated the Lys residues in the CT of both WT and CT-truncated SARS-CoV-2 S proteins and observed ∼50% reductions in particle infectivity upon overexpression of MARCH8 in the virus-producer cell. The infectivity of particles bearing truncated or Lys-mutant SARS-CoV-2 S proteins was modestly higher than those bearing the WT S protein, likely due to disruption of the endoplasmic reticulum (ER)- retention signal in the CT (48–50) (**Fig. 2D**). Expression of the inactive MARCH8-CS mutant did not reduce the infectivity of particles bearing the WT SARS-CoV-2 S protein or the full-length K-to-R mutant, but it did cause some reduction in the infectivity of virions bearing the CT-truncated forms of SARS-CoV-2 S protein (**Fig. 2D**).

We note that the effect of MARCH8 expression on the infectivity of particles bearing EboV-GP or SARS-CoV-2 S protein is modest relative to its effect on the infectivity of particles bearing VSV-G or HIV-1 Env. To investigate whether different levels of EboV-GP or SARS-CoV-2 S protein expression would affect the MARCH8-mediated inhibition of pseudotype infectivity, we performed transfections with a range of viral glycoprotein expression vector inputs. In the case of EboV-GP, we observed a 40-50% reduction in pseudotyped particle infectivity across the range of GP expression vector inputs (from 100-600 ng). In contrast, with SARS-CoV-2, we observed a 4-fold reduction in infectivity with 100 ng input, but a more-modest 2-fold reduction with a 400 ng input (data not shown).

Altogether, the data presented in Fig. 2 demonstrate that MARCH8 overexpression in the virus-producer cell inhibits the infectivity of virus particles bearing each of the four viral glycoproteins tested. The inhibitory activity of MARCH8 is largely abrogated by mutations reported to block E3 ubiquitin ligase activity (CS and W114A), although some residual inhibitory activity was observed with these mutants in some assays. Importantly, in the case of VSV-G, MARCH8-mediated inhibition is dependent on the Lys residues in the CT, whereas inhibition of HIV-1 Env, EboV-GP, and SARS-CoV-2 S protein does not require the CT.

### MARCH8 expression in virus-producer cells leads to viral envelope glycoprotein degradation and reduced processing and virion incorporation

To investigate the mechanism by which MARCH8 expression reduces the infectivity of virus particles bearing VSV-G we transfected 293T cells with the Env(-) pNL4-3 molecular clone pNL4-3/KFS and the WT or mutant VSV-G expression vectors, with or without MARCH8 expression vectors. VSV-G levels in cell and virion fractions were quantified by western blot (WB). Cotransfection with an expression vector for FLAG-tagged small glutamine-rich tetratricopeptide repeat-containing protein (SGTA) provided a loading control. As shown in **Fig. 3A**, overexpression of WT MARCH8 markedly reduced (by ∼3-fold) the expression of VSV-G in both cell and virion fractions. Consistent with the lack of inhibition observed with the MARCH8-CS and MARCH8-W114A mutants in the infectivity assays (**Fig. 2**), significant reductions in VSV-G levels were not observed with these MARCH8 mutants (**Figs. 3A and 3B**) despite comparable WT and mutant MARCH8 expression levels (**Fig. 3A**). Also consistent with the infectivity data, overexpression of WT or mutant MARCH8 did not significantly affect levels of Tailless or K-to-R VSV-G mutants in either cell or virus fractions. We also observed that overexpression of MARCH8 had no significant effect on the expression of viral Gag proteins in cells or on virus release, which, together with the observed lack of effect on SGTA expression levels, indicated that the reduction of VSV-G levels in the presence of MARCH8 is not due to cytotoxicity or globally reduced protein expression. Finally, we observed a more rapid mobility of VSV-G in the presence of WT but not mutant MARCH8, presumably due to reduced VSV-G glycosylation induced by MARCH8 expression (**Fig. 3A**). This effect on VSV-G mobility was not observed with the Tailless or K-to-R mutants, consistent with these mutants being resistant to MARCH8-mediated inhibition.

**Figure 3.**
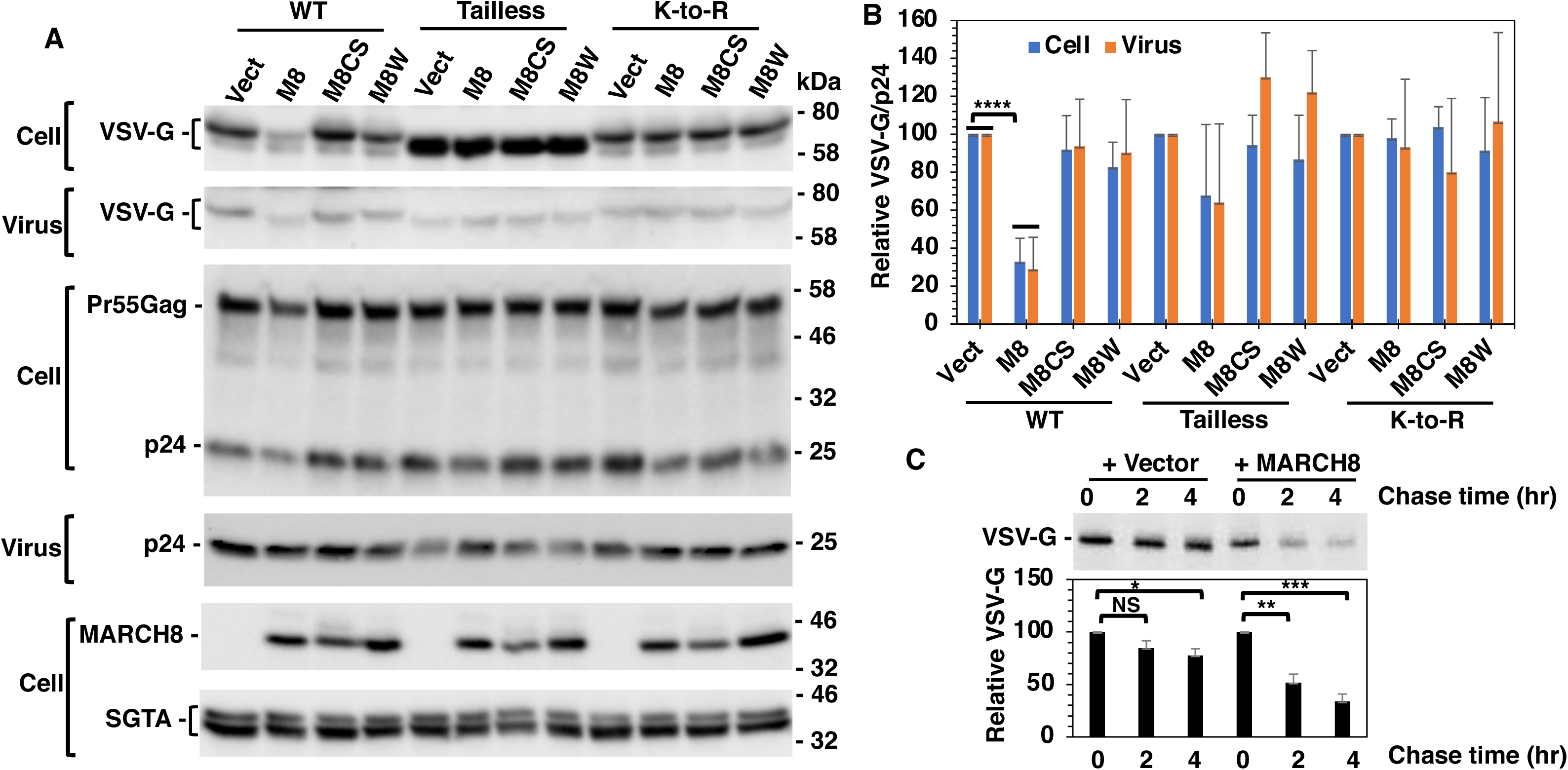
Effect of MARCH8 on virion incorporation of VSV-G is CT dependent. **(A)** HEK293T cells were cotransfected with the Env-defective HIV-1 molecular clone pNL4-3/KFS (1 μg), and vectors expressing WT or mutant VSV-G (100 ng), HA-tagged WT or mutant MARCH8 (50 ng) and FLAG-tagged SGTA (200 ng) as a transfection control. Cell and viral lysates were prepared and subjected to western blot analysis with anti-VSV-G to detect VSV-G; HIV-Ig to detect the Gag proteins Pr55Gag and p24 (CA); anti-HA to detect HA-tagged MARCH8; or anti-FLAG to detect FLAG-tagged SGTA. Mobility of molecular mass standards is shown on the right of each blot. **(B)** The levels of VSV-G in cell and virus were quantified, and the levels of VSV-G in virus were normalized to viral p24. The levels of VSV-G in the absence of MARCH8 were set to 100%. **(C)** HEK293T cells were transfected with VSV-G expression vector in the absence or presence of HA-tagged MARCH8 expression vector. One day post-transfection, cells were labeled with [^35^S]Met/Cys for 20 min and then chased for 0, 2, or 4 h in unlabeled medium. Cell lysates were prepared and immunoprecipitated with anti-VSV-G Ab and analyzed by SDS-PAGE followed by fluorography. The levels of VSV-G were quantified, and set to 100% at time zero. Data shown are ± SD from 4-5 **(B)** or 2 **(C)** independent experiments. P values (two-tailed unpaired t-test): *p < 0.05, **p < 0.02, ***p < 0.01, ****p < 0.0001.

To measure the half-life of VSV-G in the presence or absence of MARCH8, we performed pulse-chase analysis. 293T cells expressing WT VSV-G alone, or coexpressing VSV-G with MARCH8, were pulse-labeled with [^35^S]Met/Cys for 20 min and chased for 0, 2, or 4 h in unlabeled medium. We observed that MARCH8 overexpression reduced the levels of VSV-G, on average, to 52% and 34% at 2 and 4 h, respectively, compared to 85% and 78% in the absence of MARCH8 at those time points (**Fig. 3C**). These results indicate that MARCH8 overexpression promotes the degradation of VSV-G, leading to reduced expression in cells and incorporation into virions.

Similar experiments to those described above were performed with HIV-1 Env. MARCH8 expression significantly reduced the levels of both gp120 and gp41 in virions. The incorporation of WT gp120 was not impaired by expression of the MARCH8 mutants, while the MARCH8-W114A mutant modestly reduced (by ∼40%) the levels of virion-associated gp120 for CTdel144 Env. The levels of virion-associated gp41 were markedly reduced for both WT and CTdel144 Env in the presence of MARCH8, and the MARCH8 mutants did not exert a statistically significant effect on virion gp41 levels (**Fig. 4A and C**). The reduced levels of virion-associated gp120 and gp41 imposed by MARCH8 expression are consistent with the infectivity data presented above (**Fig. 2B**). The expression of exogenous FLAG-tagged SGTA and HIV-1 Gag proteins in cells was similar across the different transfected samples, indicating that expression of WT and mutant MARCH8 is not cytotoxic under these conditions and no effect on virus particle production was observed (**Fig. 4A**). These results demonstrate that MARCH8 expression in the virus-producer cell reduces the levels of virion-associated gp120 and gp41 for both WT and gp41 CT-deleted HIV-1 Env.

**Figure 4.**
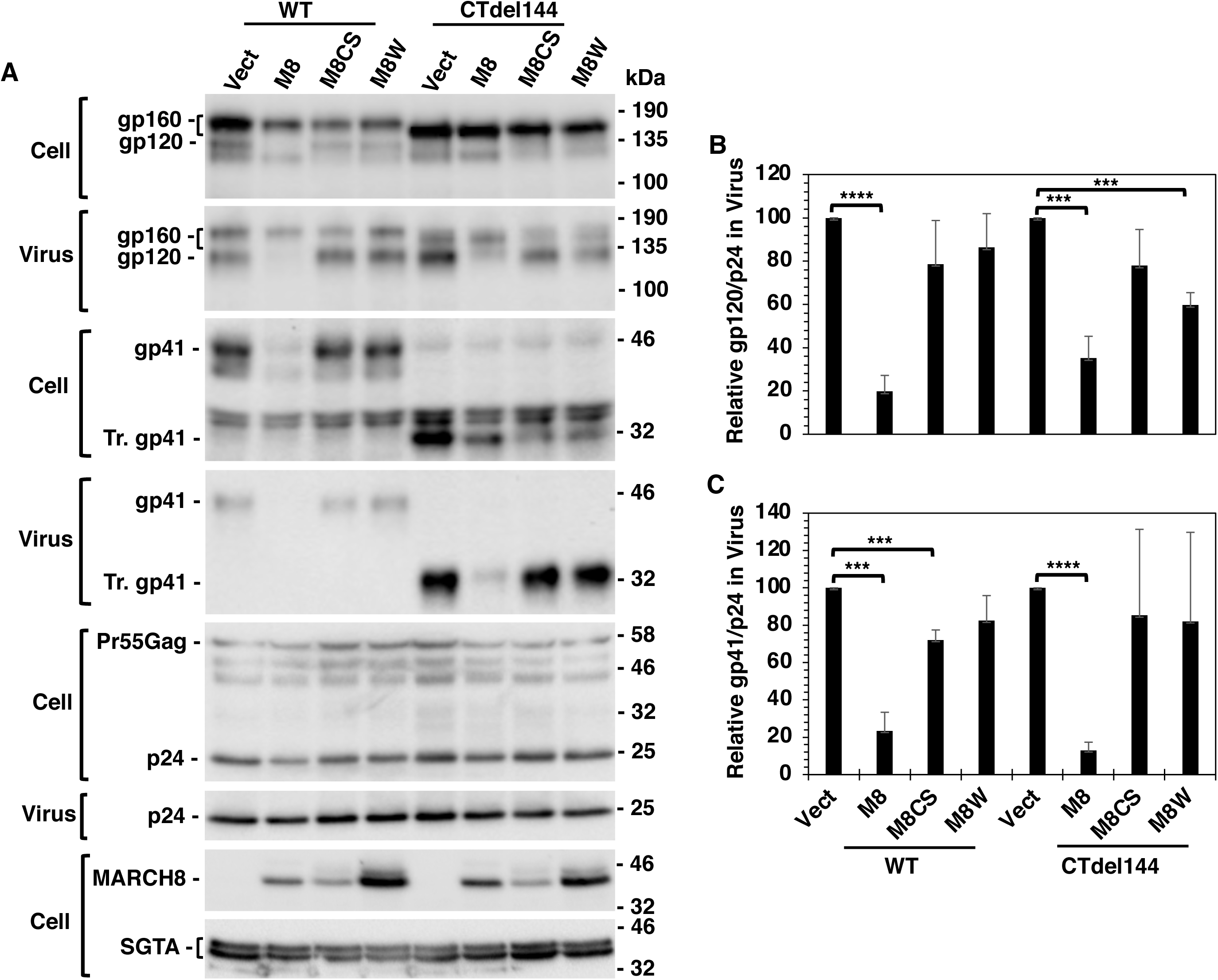
Effect of MARCH8 on HIV-1 Env processing and incorporation is CT independent. **(A)** HEK293T cells were cotransfected with pNL4-3 or pNL4-3/CTdel144 molecular clones (1 μg) in the absence or presence of vectors expressing HA-tagged WT or mutant MARCH8 (100 ng) and FLAG-tagged SGTA as a transfection control. Two days post-transfection, cell and viral lysates were prepared and subjected to western blot analysis with anti-Env to detect gp160, gp120 and gp41; HIV-Ig to detect Gag proteins, Pr55Gag and p24 (CA); anti-HA to detect HA-tagged MARCH8; or anti-FLAG to detect FLAG-tagged SGTA. Mobility of molecular mass standards is shown on the right of the blots. **(B)** The levels of gp120 in virus was quantified and normalized to p24, and set to 100% in the absence of MARCH8. **(C)** The levels of gp41 in virus was quantified and normalized to p24, and set to 100% in the absence of MARCH8. Data shown are ± SD from three independent experiments. P values (two-tailed unpaired t-test): ***p < 0.001, ****p < 0.0001.

The EboV-GP bears a CT that contains only four residues, considerably shorter than the CTs of either VSV-G or HIV-1 Env, yet it resembles HIV-1 Env in terms of the CT independence of its inhibition by MARCH8 in infectivity assays (**Fig. 2**). To investigate the effect of MARCH8 expression on the processing and incorporation of EboV-GP, we analyzed the levels of cell- and virion-associated EboV-GP in the presence and absence of MARCH8. The levels of cell- and virion-associated EboV-GP were significantly reduced in the presence of WT MARCH8, but not in the presence of the MARCH8-CS or MARCH8-W114A mutants (**Fig. 5A and C**). The incorporation of the Tailless and K-to-A EboV-GP mutants was also reduced in the presence of MARCH8 (**Fig. 5A and B**). We observed a significant reduction in the processing of EboV-GP0 in the presence of WT but not mutant MARCH8, as determined by the ratio of cell-associated GP0/GP1 (**Fig. 5A and D**). As observed with VSV-G and HIV-1 Env, overexpression of WT or mutant MARCH8 had no significant effect on cell-associated Gag expression or particle production (**Fig. 5A**). These observations demonstrate that the MARCH8-mediated infectivity defect observed with EboV-GP is due to impaired GP0 processing and correspondingly reduced virion incorporation of GP1 and GP2, and this defect is independent of the EboV-GP CT.

**Figure 5.**
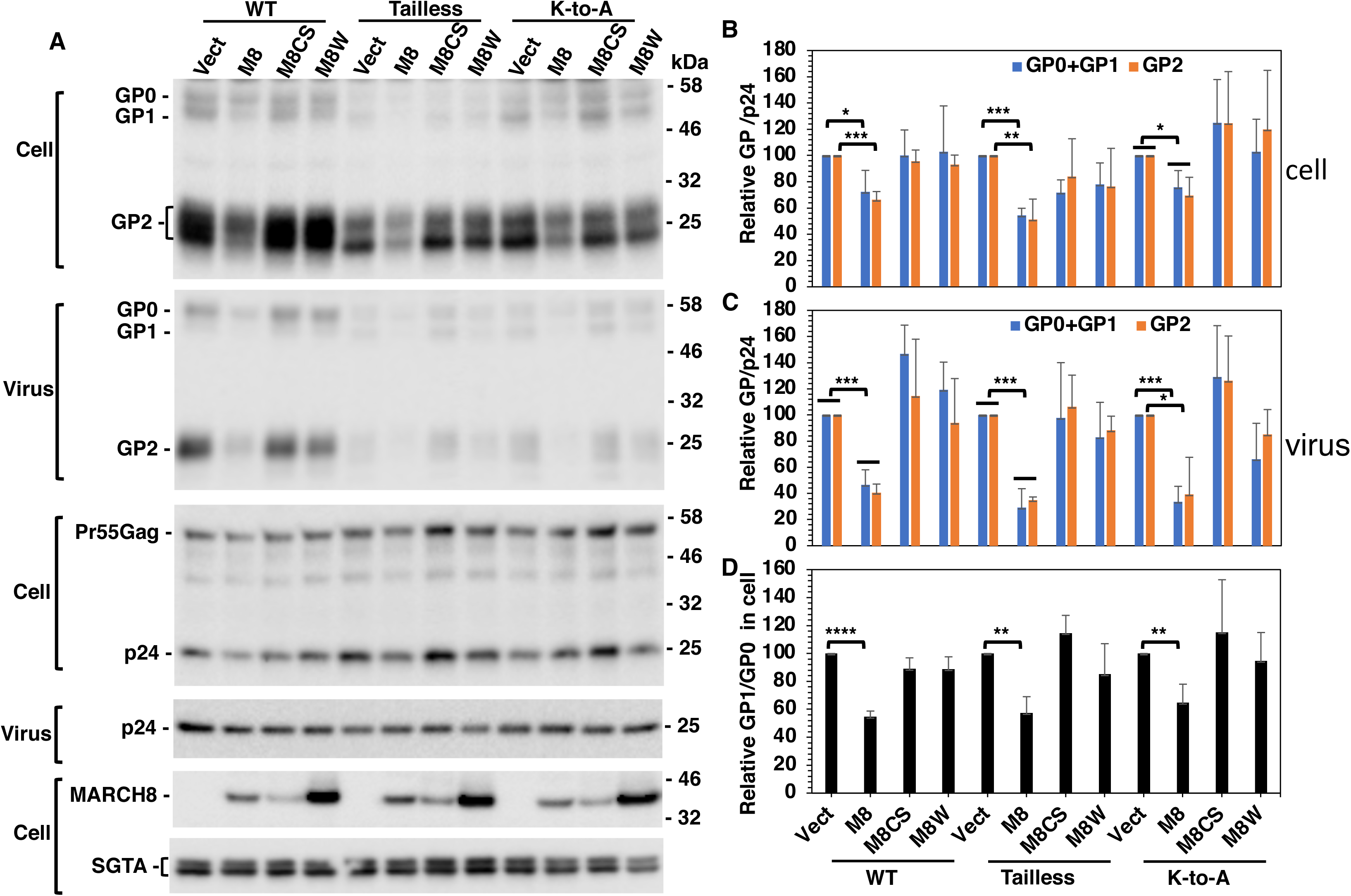
Effect of MARCH8 on processing and virion incorporation of EboV-GP is CT independent. **(A)** HEK293T cells were cotransfected with the Env-defective HIV-1 molecular clone pNL4-3/KFS (1 μg) with vectors expressing WT or mutant EboV-GP (500 ng), HA-tagged WT or mutant MARCH8 (250 ng) and FLAG-tagged SGTA as a transfection control. Two days post-transfection, cell and viral lysates were prepared and subjected to western blot analysis with anti-EboV-GP Abs to detect Pre-GP (G0), GP1, and GP2; HIV-Ig to detect Gag proteins; anti-HA to detect HA-tagged MARCH8 or anti-FLAG to detect FLAG-tagged SGTA. Mobility of molecular mass standards is shown on the right of the blots. **(B)** The levels of GP0+GP1 and GP2 in the cell were quantified, and the levels in the absence of MARCH8 were set to 100%. **(C)** The levels of GP0+GP1 and GP2 in virus were quantified and normalized to p24, and set to 100% in the absence of MARCH8. **(D)** The processing of EboV-GP in cells was quantified by calculating the ratio of GP1/GP0, and set to 100% for the no-MARCH8 control. Data shown are ± SD from three independent experiments. P values (two-tailed unpaired t-test): *p < 0.05, **p < 0.01, ***p < 0.001, ****p < 0.0001.

As mentioned in the Introduction, MARCH8 expression has been reported to be high in the lung (51), making its effect on the respiratory virus SARS-CoV-2 of particular interest. Because the experiments described above demonstrated that neither inactive MARCH8 mutant (MARCH8-CS or MARCH8-W114A) was capable of consistent antiviral activity, in these and subsequent experiments we used only one mutant, MARCH8-CS, as our negative control. As observed with HIV-1 Env and EboV-GP, the SARS-CoV-2 S protein shows a CT-independent MARCH8-mediated restriction in infectivity assays (**Fig. 2**). To examine the effect of MARCH8 on SARS-CoV-2 S protein expression, we analyzed the levels of cell- and virion-associated S protein in the presence and absence of WT or catalytically inactive MARCH8. We observed that MARCH8 expression significantly reduced (by ∼3-4 fold) the levels of S2 in cells and reduced S2 levels by ∼10-fold in virions. Averaged over multiple assays, the MARCH8-CS mutant did not significantly reduce S protein levels (**Fig. 6A and C**). Consistent with the infectivity data presented above, the levels of the S protein mutants (K-to-R, CT-truncated, and CT-truncated K-to-R) were also reduced in the presence of MARCH8. We also observed a 60-70% reduction in the levels of WT and mutant S2 in the cell fraction in the presence of MARCH8, and noted the presence of a smaller S2-related protein product (**Fig. 6A, red arrow**), which we speculate to be a less-glycosylated form of S2. The reductions in S2, and the presence of the smaller S2 species, were not observed upon expression of the MARCH8-CS mutant (**Fig. 6A and B**). Levels of the S precursor protein were not reduced by expression of MARCH8 in the virus-producer cell, suggesting an effect primarily on S protein processing. These data demonstrate that expression of WT MARCH8 but not the MARCH8-CS mutant disrupts S protein processing, glycosylation, and incorporation into virus particles and indicate that the MARCH8-imposed antagonism of SARS-CoV-2 S protein is CT independent.

**Figure 6.**
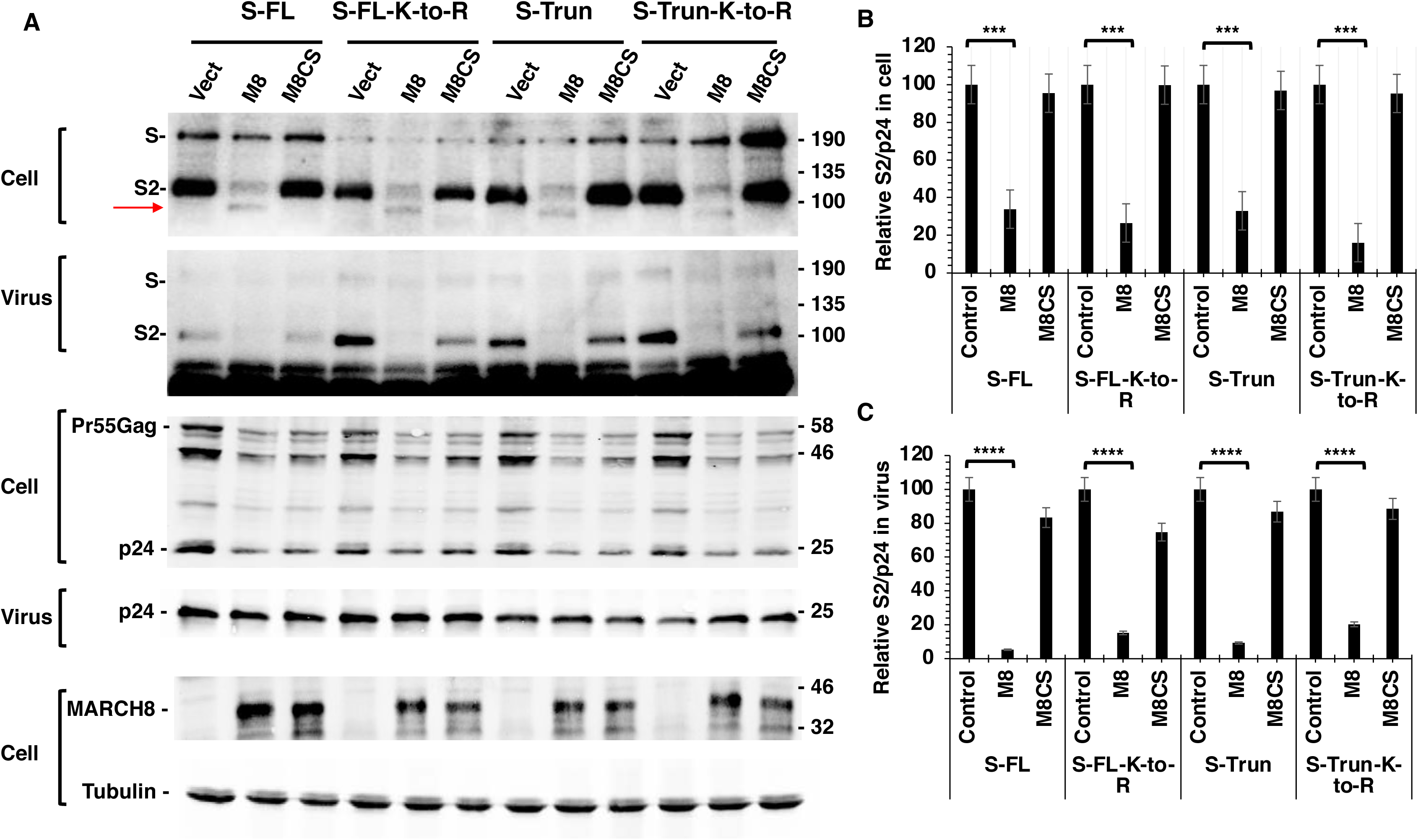
Effect of MARCH8 on processing and virion incorporation of SARS-CoV-2-S is CT independent. **(A)** HEK293T were cotransfected with the Env-defective, luciferase-expressing pNL4-3 derivative pNL4-3.Luc.R-E- (3 μg) with vectors expressing WT or mutant SARS-CoV-2 S protein (300 ng), and WT or mutant HA-tagged MARCH8 (200 ng). Two days post-transfection, cell and virus lysate were prepared and subjected to western blot analysis with anti-SARS-CoV-2 S2 Ab to detect precursor S and S2; HIV-Ig to detect Gag proteins, Pr55Gag and p24; and anti-HA to detect HA-tagged MARCH8. Molecular mass standards are shown on the right of the blots. Red arrow indicates a putative less-glycosylated form SARS-CoV-2 S2 protein in the cell lysate. **(B)** The level of SARS-CoV-2 S protein in cells was quantified and normalized to p24; values were set to 100% in the absence of MARCH8. **(C)** The level of S2 was quantified and normalized to p24 in virus; values were set to 100% in the absence of MARCH8. Data shown are ± SD from three independent experiments. P values (two-tailed unpaired t-test): ***p < 0.001, ****p < 0.0001.

Because the effect of MARCH8 expression on both particle infectivity and glycoprotein levels in virions varied across the four viral glycoproteins studied here, we calculated the incorporation efficiency of each glycoprotein under the conditions used in the experiments described above (**Fig. S2**). We observed that the Ebo-GP incorporation efficiency was quite high, with nearly 40% of expressed GP present in VLPs. In contrast, the incorporation of SARS-CoV-2 S protein was quite low, perhaps because HIV-1 particles assemble on the plasma membrane, which, as noted in the Introduction, is not the normal site of SARS-CoV-2 assembly. Incorporation efficiencies of HIV-1 gp41 and VSV-G were ∼18% and ∼8%, respectively, under these experimental conditions (**Fig. S2**).

### Knockdown of endogenous MARCH8 expression in HEK293T increases HIV-1 infectivity

Although levels of basal MARCH8 expression are known to be relatively low in established cell lines (51), we examined whether knock-down of *MARCH8* gene expression in HEK293T cells would affect the infectivity of HIV-1 particles produced from the *MARCH8*- depleted cells. HEK293T cells were treated with siRNA targeting MARCH8 or with a non-targeting control. *MARCH8* expression was measured by RT-qPCR. At the highest siRNA input, *MARCH8* gene expression was reduced by approximately 5-fold (**Fig. 7A**). Under these conditions, we observed a modest but statistically significant increase in the infectivity of HIV-1 particles produced from the *MARCH8*-depleted cells (**Fig. 7B**). The infectivity of virus produced from cells treated with the non-targeting siRNA control was not affected. These results demonstrate that even low levels of endogenous MARCH8 expression can impair HIV-1 infectivity.

**Figure 7.**
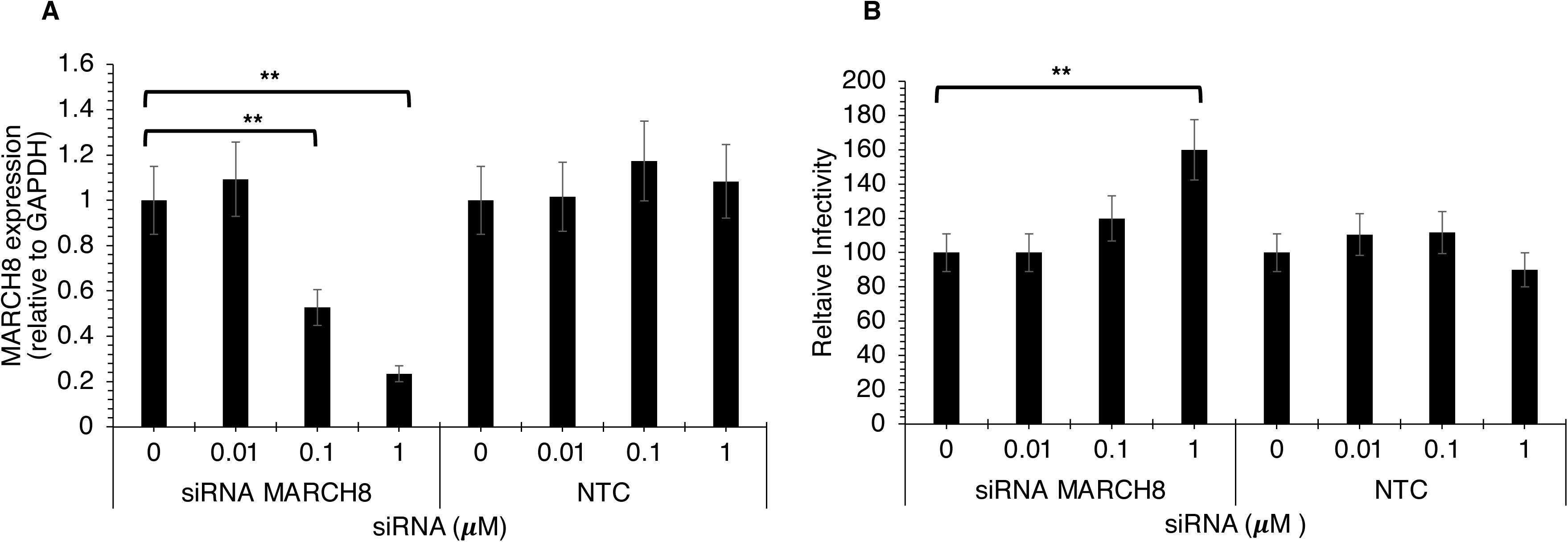
Knockdown of endogenous MARCH8 in HEK293T cells increases HIV-1 infectivity. A) HEK293T cells were treated for 72 hrs with the indicated concentrations of MARCH8-specific siRNA or a non-targeting control (NTC). Endogenous *MARCH8* RNA expression was measured using RT-qPCR. B) siRNA-treated HEK293T cells in a 6-well plate were transfected with the pNL4-3 HIV-1 molecular clone (3 μg). Two days post-transfection, virus supernatants were collected, RT normalized, and used to infect TZM-bl cells. Infectivity in non-siRNA-treated cells is set to 100%. Data shown are ± SD from 3 independent experiments. P values (two-tailed unpaired t-test): *p < 0.05, **p < 0.01.

### *MARCH8* expression is highly and rapidly inducible in certain cell types by IFN stimulation

Previous findings demonstrated that MARCH8 is highly expressed in MDMs in the absence of IFN stimulation and that expression levels are not appreciably increased in either MDMs or human CD4^+^ T cells by overnight stimulation with IFN-α (11). Because IFN is known to induce highly variable levels of gene expression in a time-, dose-, and cell type-dependent manner (52, 53), and many genes with antiviral functions are IFN inducible (54), we performed RT-qPCR time-course experiments with three different IFNs [IFN type-I (α and β) and II (γ)] using total RNA extracted from different cell types and cell lines. We tested the SupT1 human T-cell line, primary hPBMCs from four different donors, the A549 transformed alveolar epithelial cell line, primary human airway epithelial cells, and HEK293T cells. In the SupT1 T-cell line, *MARCH8* gene expression was rapidly induced at 0.5 hr post-stimulation with IFN-α up to 70- fold over baseline levels, which subsided to ∼10-fold over unstimulated levels by 4 hrs after induction (**Fig. 8A**) and then returned to baseline. Treatment with IFN-β induced a ∼30-fold increase in *MARCH8* gene expression over baseline that peaked around 1-2 hrs post-IFN stimulation (**Fig. 8A**). To examine *MARCH8* gene expression in primary hPBMCs, cells from four donors were IFN stimulated and *MARCH8* RNA levels measured. In three of the four donors, we observed that *MARCH8* expression increased 5-8-fold over unstimulated levels at 0.5 hr post-stimulation with IFN-α and then returned to baseline by 2 hr post-stimulation (**Fig. 8B**). In one donor, no IFN-inducible *MARCH8* gene expression was observed (data not shown). To investigate *MARCH8* gene expression in cells that are permissive for SARS-CoV-2 infection, we tested primary human airway epithelial cells and observed a ∼3-fold increase in expression at 1-4 h post IFN-α stimulation (**Fig. 8C**). In HEK293T cells, we measured a rapid, ∼7-fold increase in MARCH8 gene expression after stimulation with IFN-α. Finally, we observed no IFN stimulation of *MARCH8* gene expression in A549 cells (data not shown). Collectively, these data indicate that *MARCH8* gene expression is induced to varying levels in cell lines and primary cells types that are natural targets for HIV-1 and SARS-CoV-2 infection.

**Figure 8.**
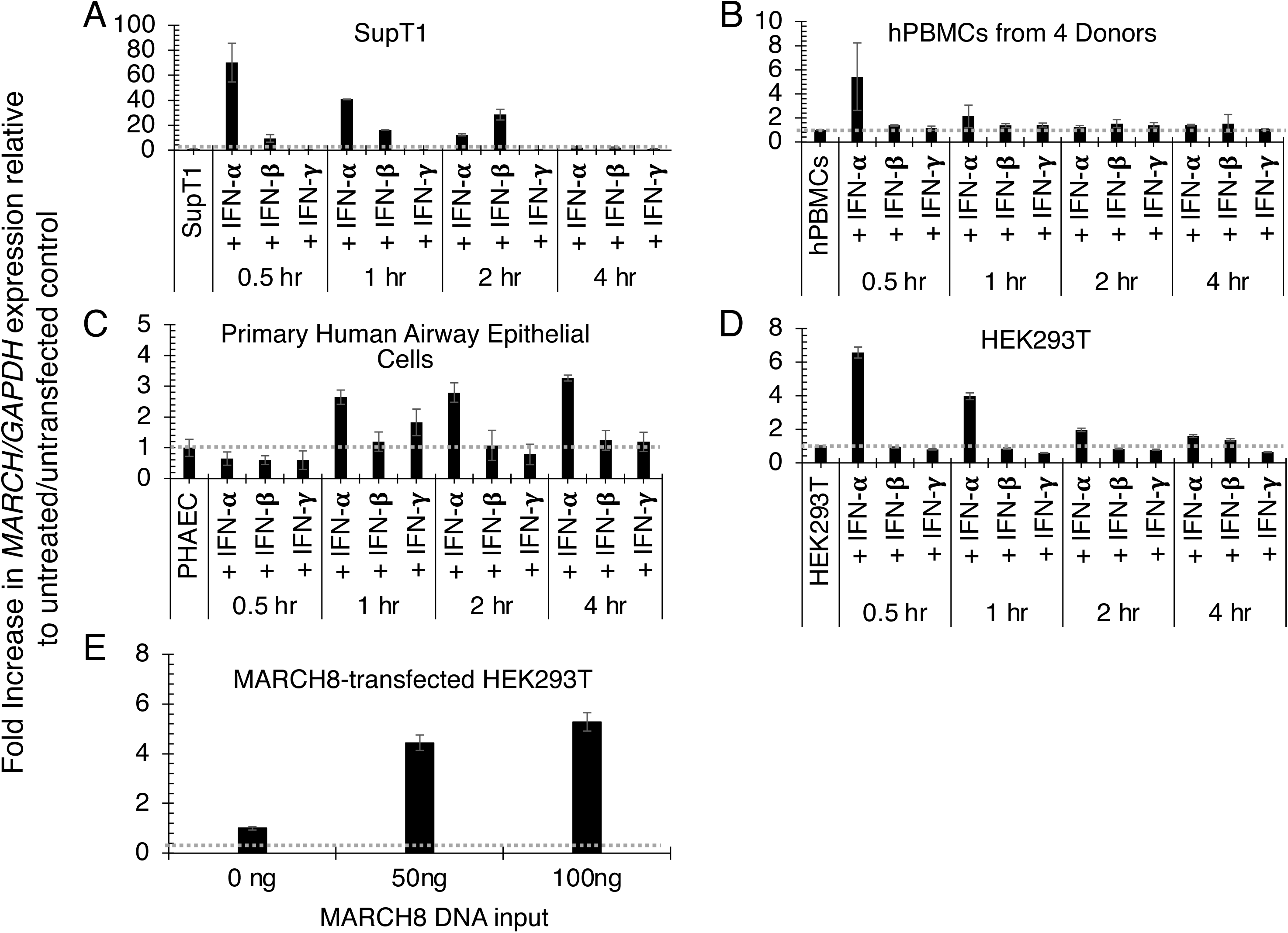
Endogenous *MARCH8* gene expression is rapidly induced by IFN. Endogenous *MARCH8* RNA expression in the SupT1 T-cell line **(A)**, hPBMCs from 4 donors **(B),** primary human airway epithelial cells **(C),** and the HEK293T cell line **(D)** at 0, 0.5, 1, 2, and 4 hrs post-stimulation with 1000U/ml type I (α and β) and II (γ) IFN. **E**) *MARCH8* expression in transfected HEK293T cells 48 hr post-transfection (hpt) with increasing concentrations of MARCH8 expression vector (0, 50, 100 ng) in a 6-well tissue culture plate. Isolated RNA was subjected to the RT-qPCR protocol as detailed in the Materials and Methods. Fold increase in *MARCH8* RNA expression relative to untreated (A-D) or untransfected (E) controls is indicated. *MARCH8* RNA levels are expressed as a ratio of *MARCH8*/*GAPDH*. Data shown are ± SD from three independent experiments.

It is important to verify that the levels of exogenous *MARCH8* gene expression achieved in these assays were not excessive relative to endogenous levels present in relevant cell types. Therefore, we investigated the *MARCH8* RNA expression levels in transiently transfected HEK293T cells upon increasing MARCH8 concentrations at 48 hr post-transfection. The results indicated that *MARCH8* expression at 0.05 and 0.1 μg DNA input, the conditions used for the analyses presented in Figs. 2-6, was ∼4-5 fold above the levels in untransfected cells (**Fig. 8E**). Transfection efficiencies were ∼50% under these conditions (data not shown), indicating that transfected cells expressed *MARCH8* RNA at levels that were ∼8-10 fold over those of untransfected cells. These results show that exogenous levels of *MARCH8* gene expression in our transfected 293T cells were lower than those observed following IFN stimulation of the SupT1 T-cell line and comparable to those measured in some IFN-stimulated hPBMC donors and HEK293T cells at early time points after IFN stimulation (**Fig. 8A-D**). The levels of basal expression of *MARCH8* relative to the house-keeping gene *GAPDH* in all of these cell types were comparable (**Table S1**). Collectively, our data demonstrate that *MARCH8* gene expression can be induced rapidly by IFN stimulation in different cell types: a human T-cell line, hPBMCs, human airway epithelial cells and the HEK293T cell line. The levels of exogenous expression in our transfected HEK293T cells were lower than, or comparable to, those achieved upon IFN stimulation at early time points in the T-cell line, primary hPBMCs, and HEK293T cells.

### MARCH8 retains VSV-G, HIV-1 Env, EboV-GP and SARS-CoV-2 S protein in an intracellular compartment

The western blotting data presented above demonstrate that MARCH8 overexpression disrupts the processing and incorporation of viral envelope glycoproteins, suggesting that MARCH8 might alter the trafficking of these viral glycoproteins. To examine the effect of MARCH8 overexpression on the localization of different viral glycoproteins, we performed confocal microscopy using HEK293T cells transfected with MARCH8 or the inactive MARCH8-CS mutant. Our confocal data showed that MARCH8 localizes to the Golgi, a LAMP-1^+^ compartment, and the plasma membrane (**Fig. S1**) as previously reported (51). All of the viral glycoproteins and mutants used in this study colocalized with MARCH8 and MARCH8-CS with strong Pearson’s correlation coefficients (r) of ∼0.7-0.9 (Table 1) indicating that MARCH proteins and viral glycoproteins traffic through the same pathway. We quantified the intracellular colocalization between the dsRed-Golgi marker and the different viral glycoproteins in the presence of WT MARCH8 and the inactive MARCH8-CS and observed strong colocalization (r=∼0.7-0.9) (**Table 1, Fig. 9 to 12**).

**Figure 9.**
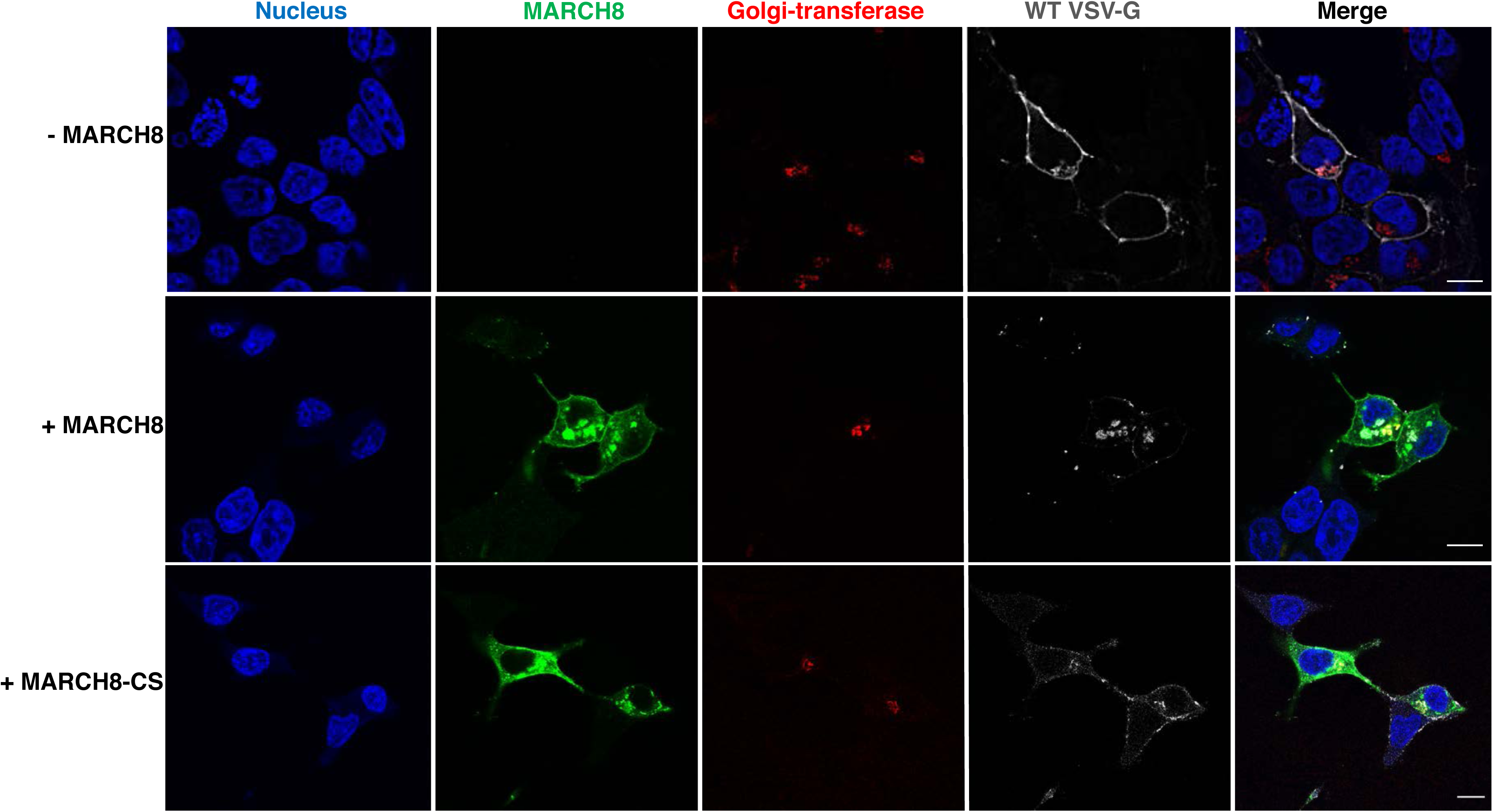
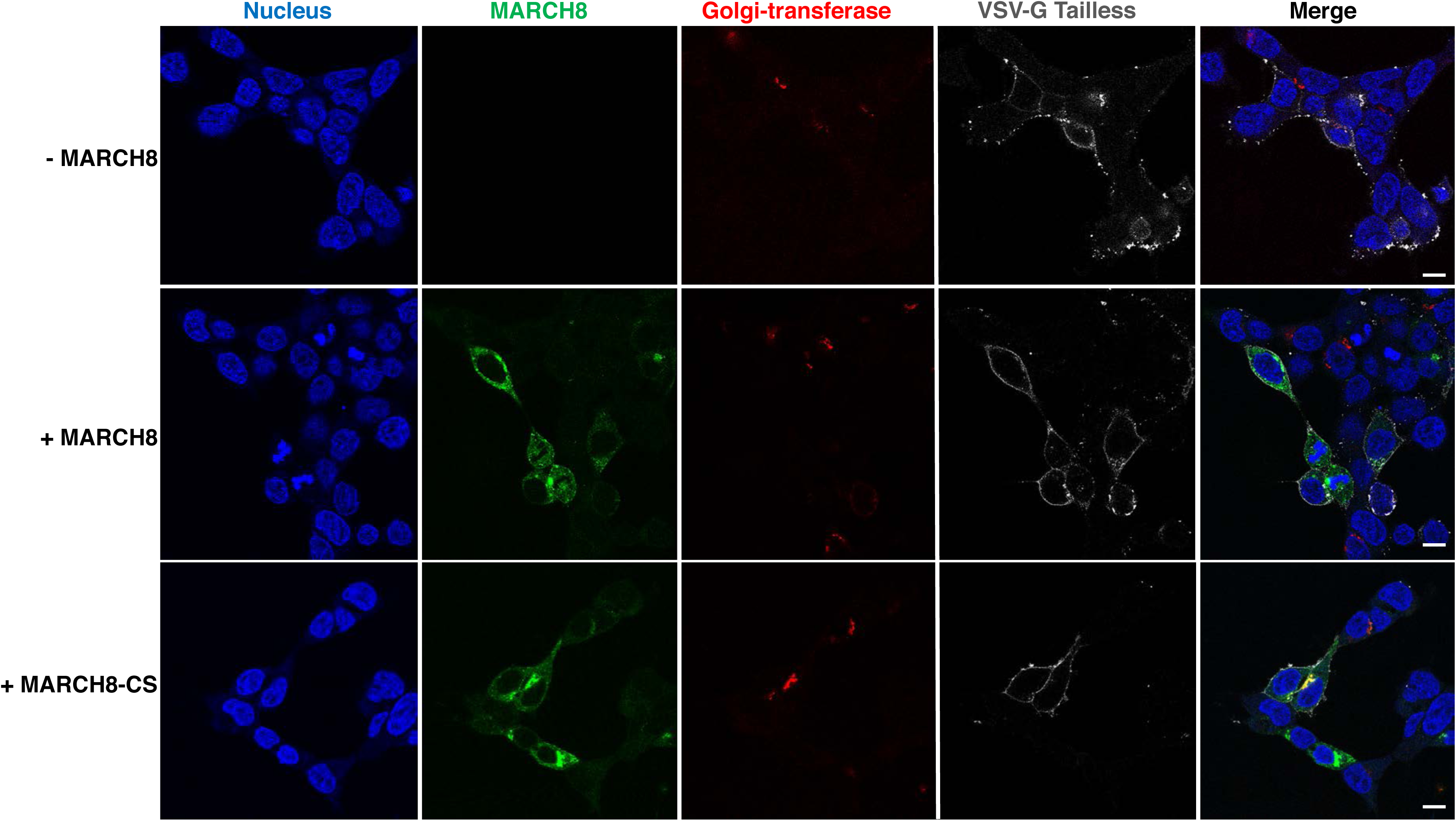
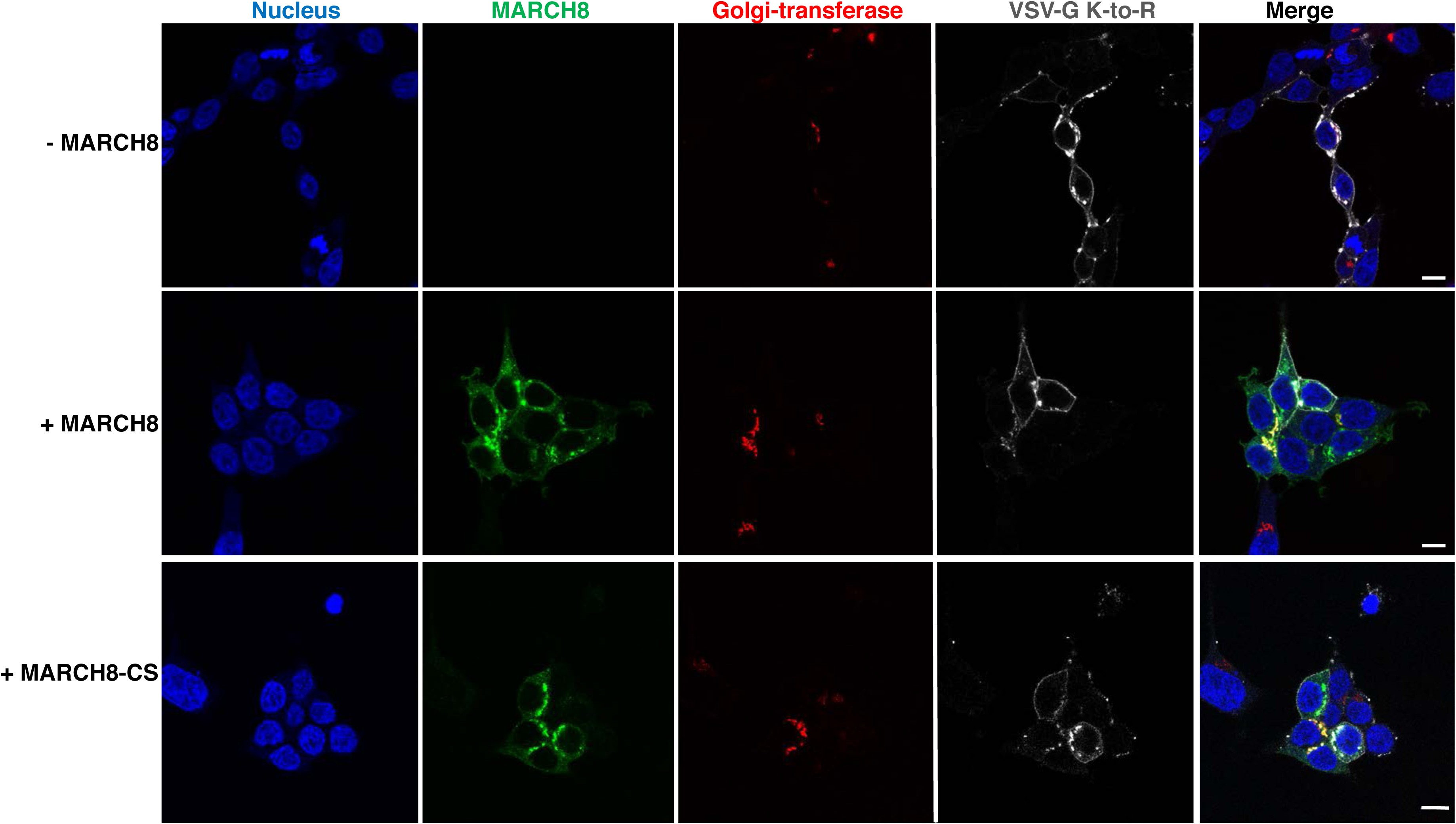
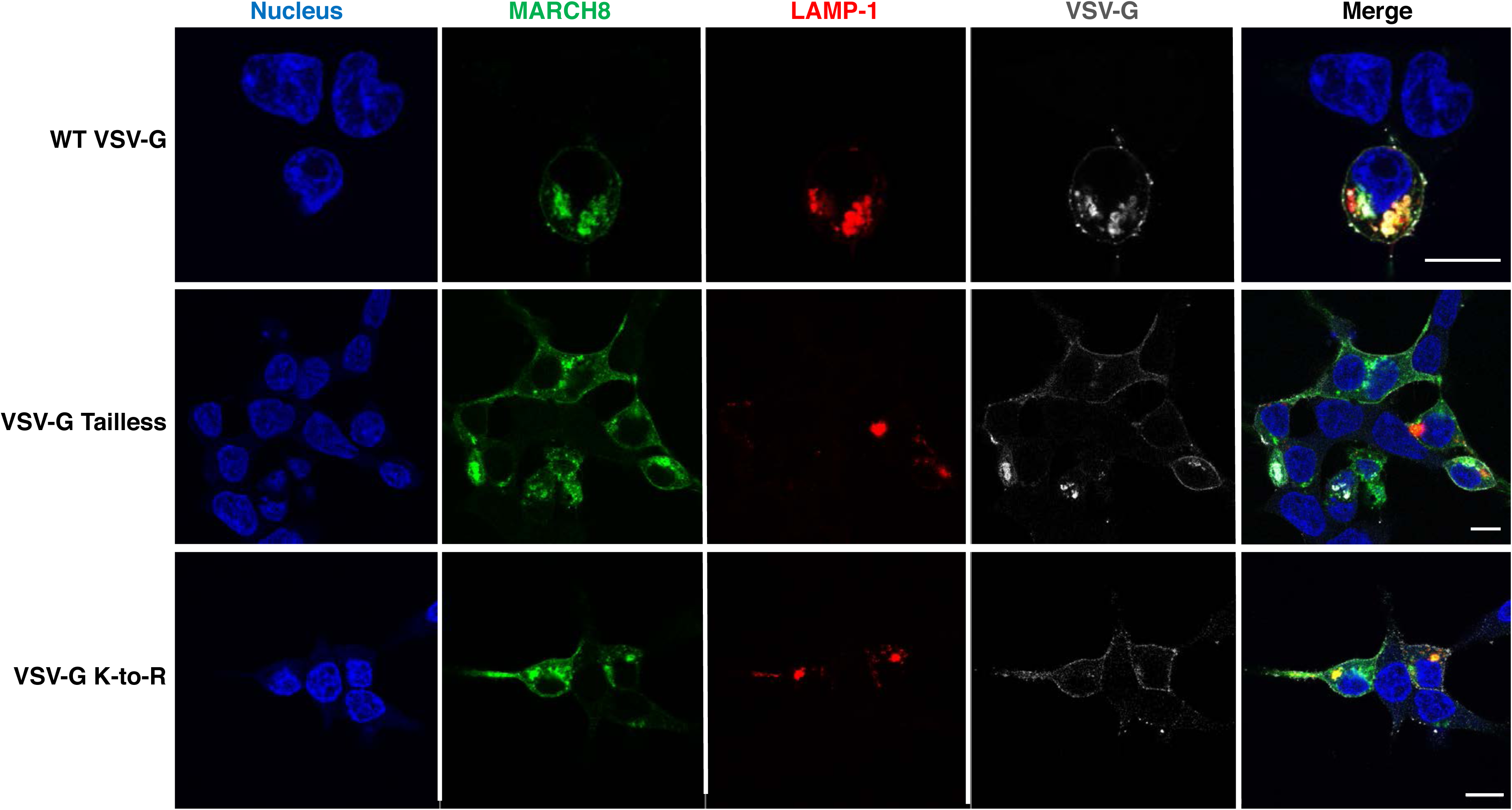
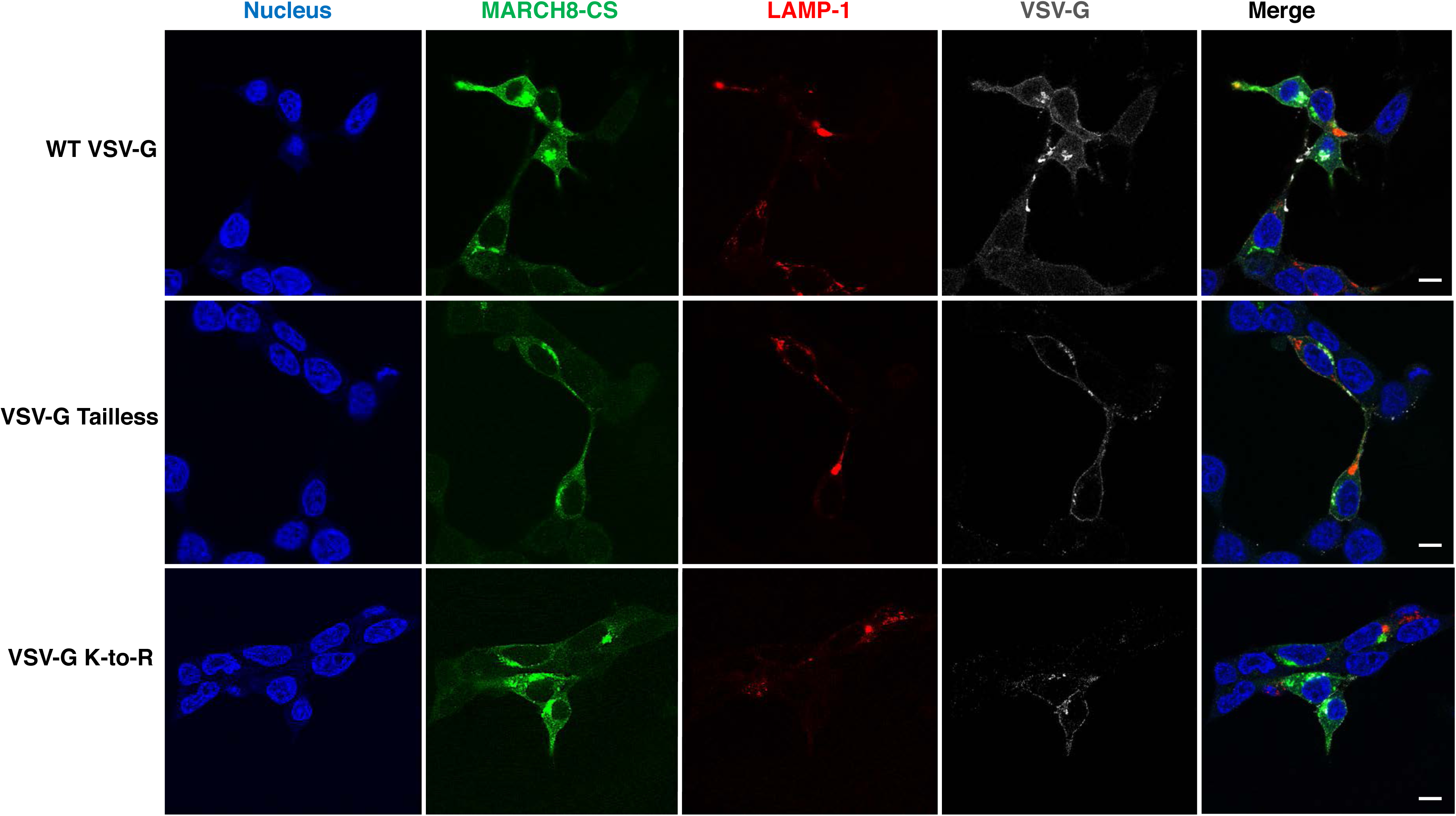
MARCH8 traps WT VSV-G but not the VSV-G CT mutants in an intracellular LAMP-1^+^ compartment. HEK293T cells were co-transfected with vectors expressing WT or mutant VSV-G (200 ng) with or without vectors expressing HA-tagged MARCH8, MARCH8-CS (100 ng) and the LAMP1-RFP or pDsRed-Golgi-Beta 1,4-galactosyltransferase (Golgi-transferase) (300 ng) expression vectors. One day post-transfection, cells were processed for confocal microscopy. Distribution of WT VSV-G **(A)**, VSV-G Tailless **(B)**, or VSV-G K-to-R **(C)** in the absence or presence of WT MARCH8 or MARCH8-CS. Colocalization of VSV-G or VSV-G mutants with WT MARCH8 **(D)** or MARCH8-CS **(E)** with the LAMP1-RFP or pDsRed-Golgi-Beta 1,4-galactosyltransferase cellular markers. Scale bar = 10 µm.

**Table 1.**
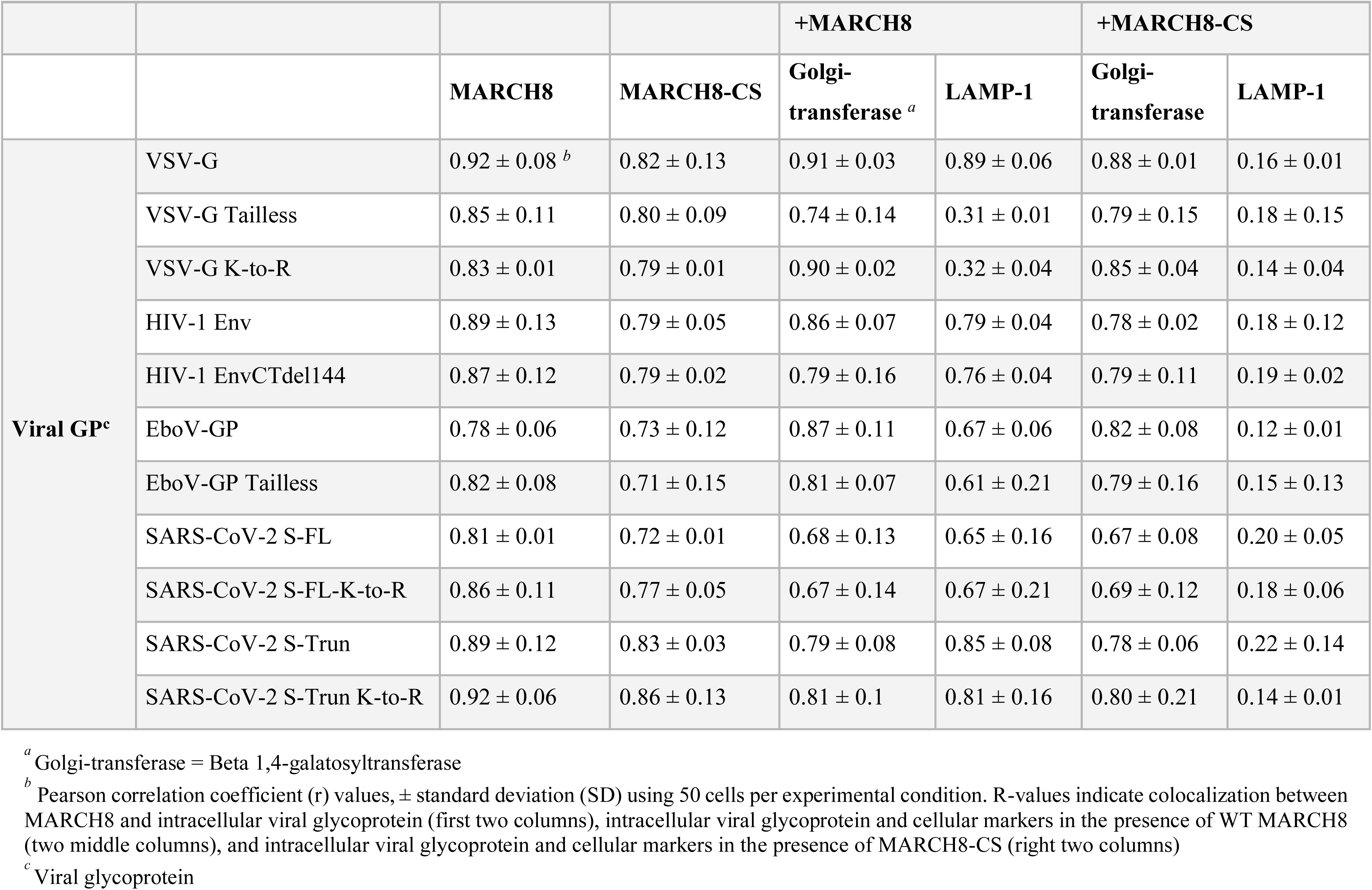
Colocalization of WT and mutant MARCH8, viral glycoproteins, and cellular markers

|  |  |  |  |  |  |  |  |
| --- | --- | --- | --- | --- | --- | --- | --- |
|  |  |  |  | +MARCH8 |  | +MARCH8-CS |  |
|  |  | MARCH8 | MARCH8-CS | Golgi-transferase <sup>a</sup> | LAMP-1 | Golgi-transferase | LAMP-1 |
| Viral GP<sup>c</sup> | VSV-G | 0.92 ± 0.08 <sup>b</sup> | 0.82 ± 0.13 | 0.91 ± 0.03 | 0.89 ± 0.06 | 0.88 ± 0.01 | 0.16 ± 0.01 |
|  | VSV-G Tailless | 0.85 ± 0.11 | 0.80 ± 0.09 | 0.74 ± 0.14 | 0.31 ± 0.01 | 0.79 ± 0.15 | 0.18 ± 0.15 |
|  | VSV-G K-to-R | 0.83 ± 0.01 | 0.79 ± 0.01 | 0.90 ± 0.02 | 0.32 ± 0.04 | 0.85 ± 0.04 | 0.14 ± 0.04 |
|  | HIV-1 Env | 0.89 ± 0.13 | 0.79 ± 0.05 | 0.86 ± 0.07 | 0.79 ± 0.04 | 0.78 ± 0.02 | 0.18 ± 0.12 |
|  | HIV-1 EnvCTdel144 | 0.87 ± 0.12 | 0.79 ± 0.02 | 0.79 ± 0.16 | 0.76 ± 0.04 | 0.79 ± 0.11 | 0.19 ± 0.02 |
|  | EboV-GP | 0.78 ± 0.06 | 0.73 ± 0.12 | 0.87 ± 0.11 | 0.67 ± 0.06 | 0.82 ± 0.08 | 0.12 ± 0.01 |
|  | EboV-GP Tailless | 0.82 ± 0.08 | 0.71 ± 0.15 | 0.81 ± 0.07 | 0.61 ± 0.21 | 0.79 ± 0.16 | 0.15 ± 0.13 |
|  | SARS-CoV-2 S-FL | 0.81 ± 0.01 | 0.72 ± 0.01 | 0.68 ± 0.13 | 0.65 ± 0.16 | 0.67 ± 0.08 | 0.20 ± 0.05 |
|  | SARS-CoV-2 S-FL-K-to-R | 0.86 ± 0.11 | 0.77 ± 0.05 | 0.67 ± 0.14 | 0.67 ± 0.21 | 0.69 ± 0.12 | 0.18 ± 0.06 |
|  | SARS-CoV-2 S-Trun | 0.89 ± 0.12 | 0.83 ± 0.03 | 0.79 ± 0.08 | 0.85 ± 0.08 | 0.78 ± 0.06 | 0.22 ± 0.14 |
|  | SARS-CoV-2 S-Trun K-to-R | 0.92 ± 0.06 | 0.86 ± 0.13 | 0.81 ± 0.1 | 0.81 ± 0.16 | 0.80 ± 0.21 | 0.14 ± 0.01 |

<sup>a</sup> Golgi-transferase = Beta 1,4-galatosyltransferase
<sup>b</sup> Pearson correlation coefficient (r) values, ± standard deviation (SD) using 50 cells per experimental condition. R-values indicate colocalization between MARCH8 and intracellular viral glycoprotein (first two columns), intracellular viral glycoprotein and cellular markers in the presence of WT MARCH8 (two middle columns), and intracellular viral glycoprotein and cellular markers in the presence of MARCH8-CS (right two columns)
<sup>c</sup> Viral glycoprotein

In the absence of MARCH8 overexpression, we observed WT VSV-G localization primarily in the Golgi and at the plasma membrane (**Fig. 9A**). MARCH8 but not MARCH8-CS expression resulted in a marked shift in the localization of WT VSV-G from the plasma membrane to an intracellular vesicular compartment (**Fig. 9A**). The predominantly plasma membrane localization of the VSV-G Tailless (**Fig. 9B**) and VSV-G K-to-R (**Fig. 9C**) mutants was not shifted by MARCH8 expression, consistent with the lack of MARCH8 targeting of these VSV-G mutants in infectivity and western blotting analyses. We observed a strong colocalization (r ∼ 0.89) between WT VSV-G and the lysosomal marker LAMP-1 in the presence of WT MARCH8 expression (**Fig. 9D; Table 1**), whereas the Tailless and K-to-R VSV-G mutants showed low levels of colocalization with LAMP-1 (r ∼ 0.2). In the presence of the inactive MARCH8-CS mutant, neither WT nor mutant VSV-G showed a high level of colocalization with LAMP-1 (r ∼ 0.2) (**Fig. 9E; Table 1**). These data demonstrate that MARCH8 interferes with VSV-G trafficking and targets it to a LAMP-1-positive compartment in a manner that requires the Lys residues in the VSV-G CT.

We performed similar analyses with HIV-1 Env. As expected, WT and CTdel144 HIV-1 Env were observed in the Golgi and at the plasma membrane in the absence of MARCH8 expression (**Fig. 10A and B**). Similar to VSV-G, localization of WT HIV-1 Env was shifted in cells expressing MARCH8, and a high level of colocalization (r ∼0.8) with LAMP-1 was observed (**Fig. 10C; Table 1**). In contrast to VSV-G, for which MARCH8-induced localization required the CT, the Tailless form of HIV-1 Env, CTdel144, was strongly localized to a LAMP-1^+^ compartment in MARCH8-expressing cells (**Fig. 10C; Table 1**). The inactive MARCH8-CS did not result in the retention of either WT or CTdel144 HIV-1 Env in a LAMP-1^+^ compartment (**Fig. 10D**). These findings are consistent with the infectivity and western blotting data presented above, and indicate that the reductions in HIV-1 Env processing and incorporation in the presence of MARCH8 are associated with the retention of Env in an internal, LAMP-1^+^ compartment.

**Figure 10.**
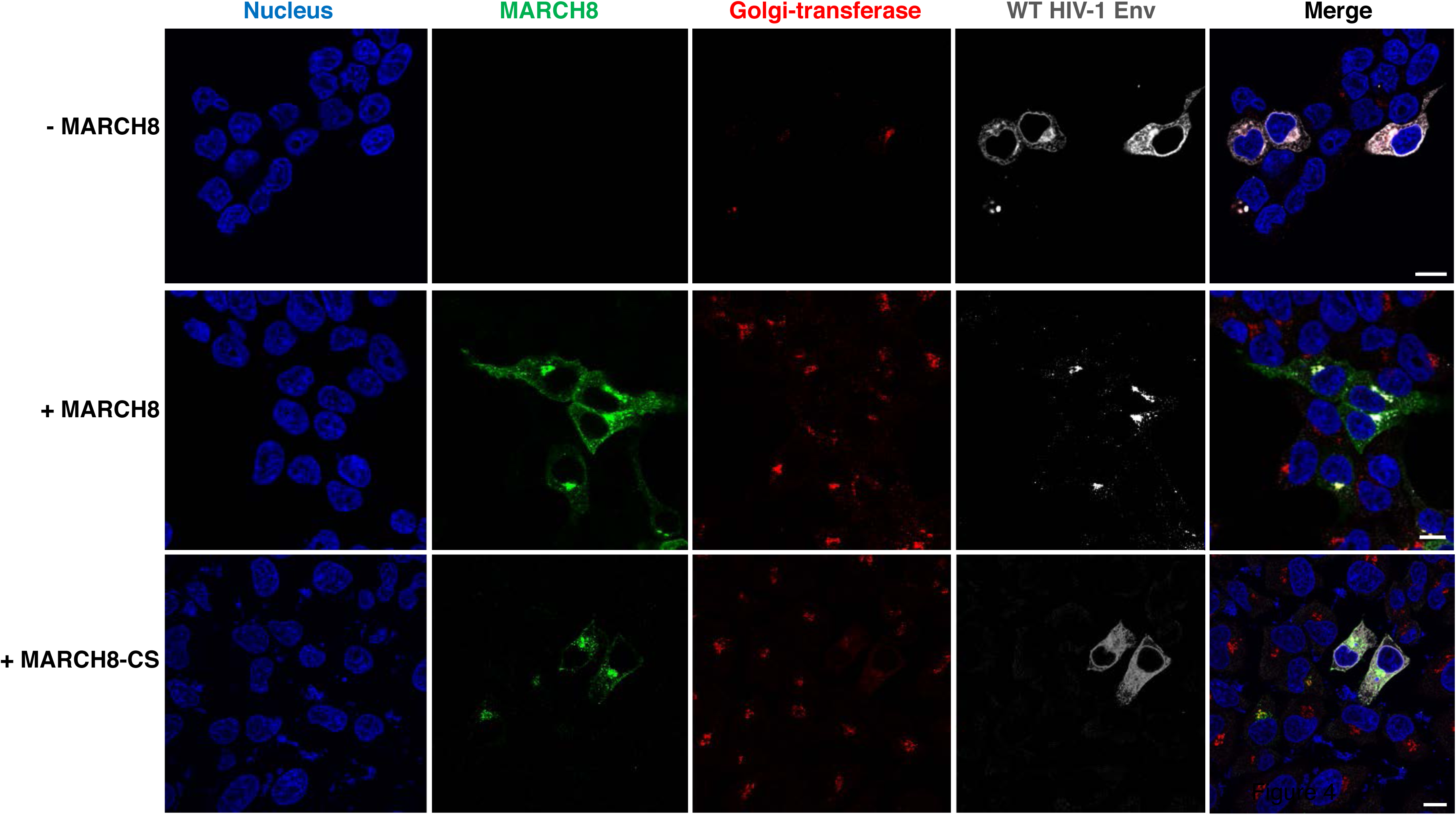
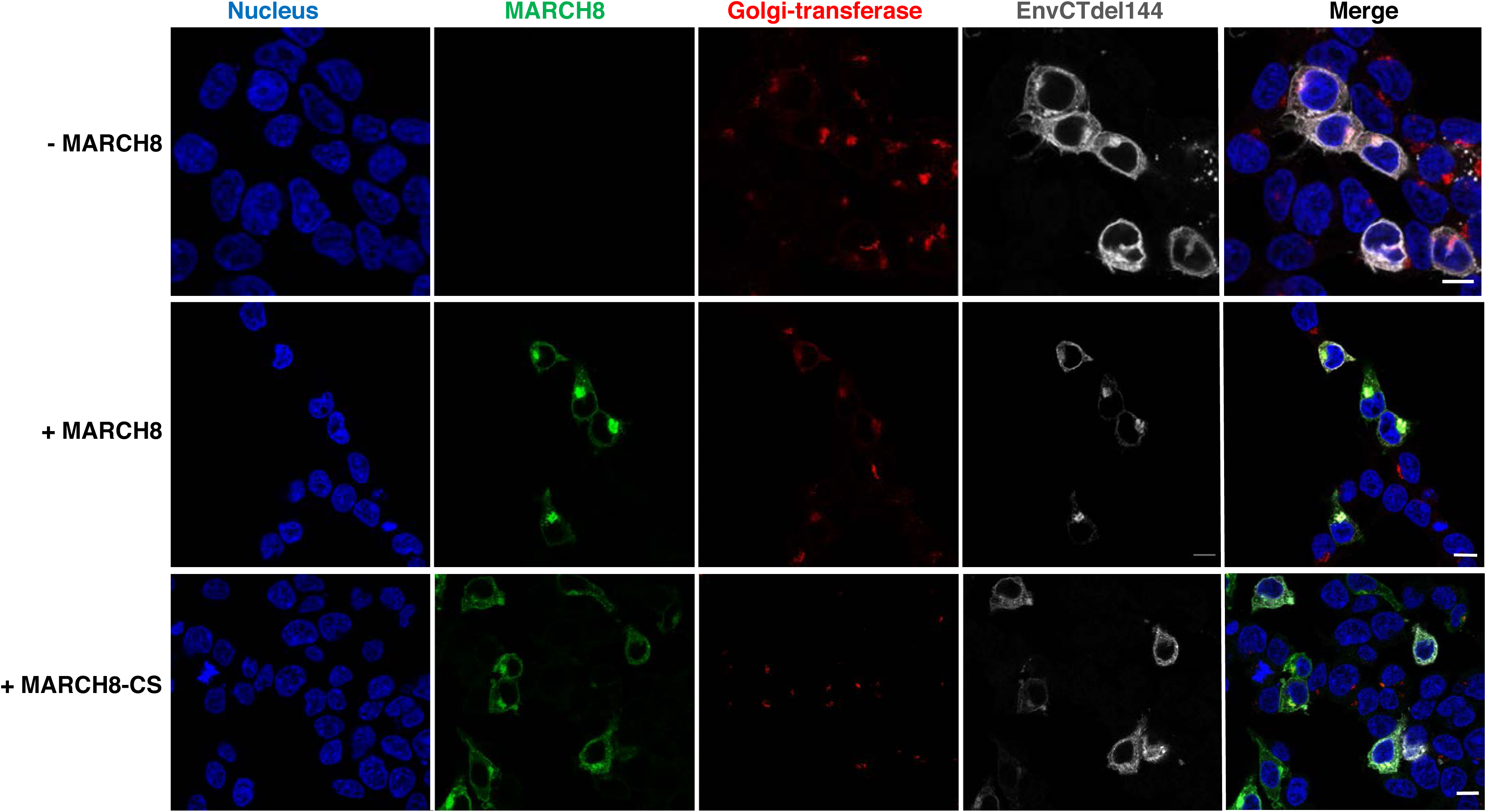
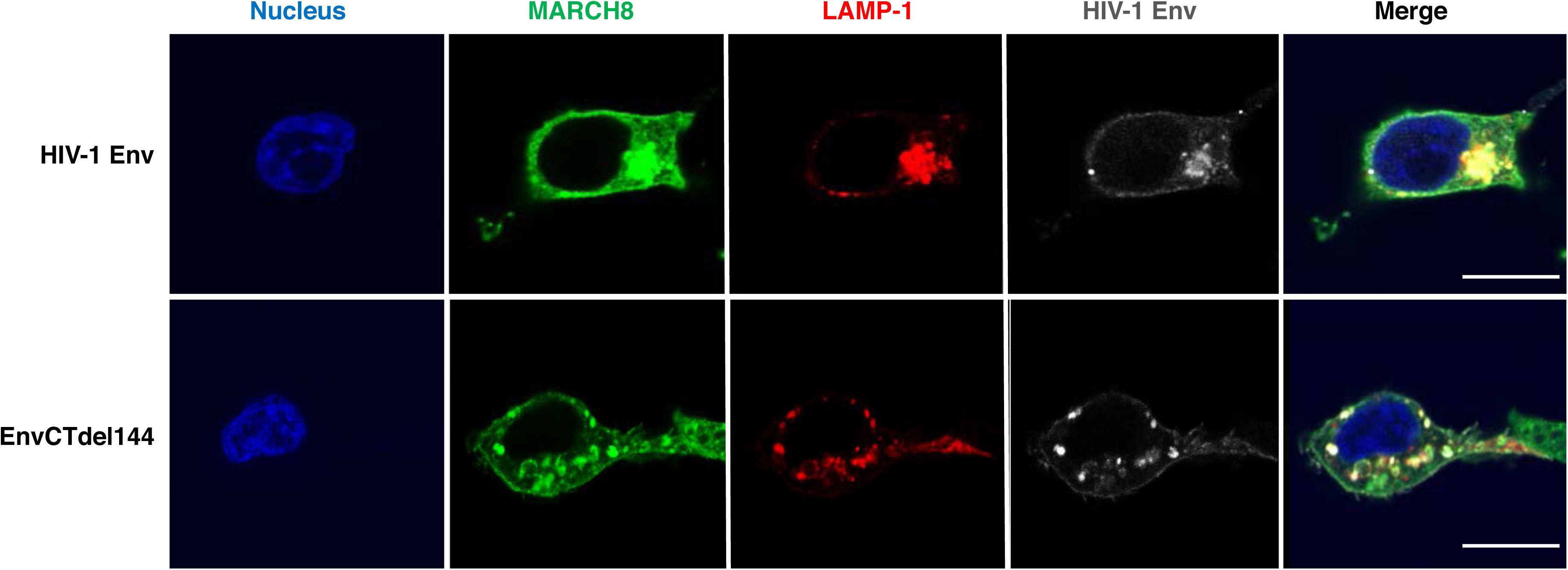
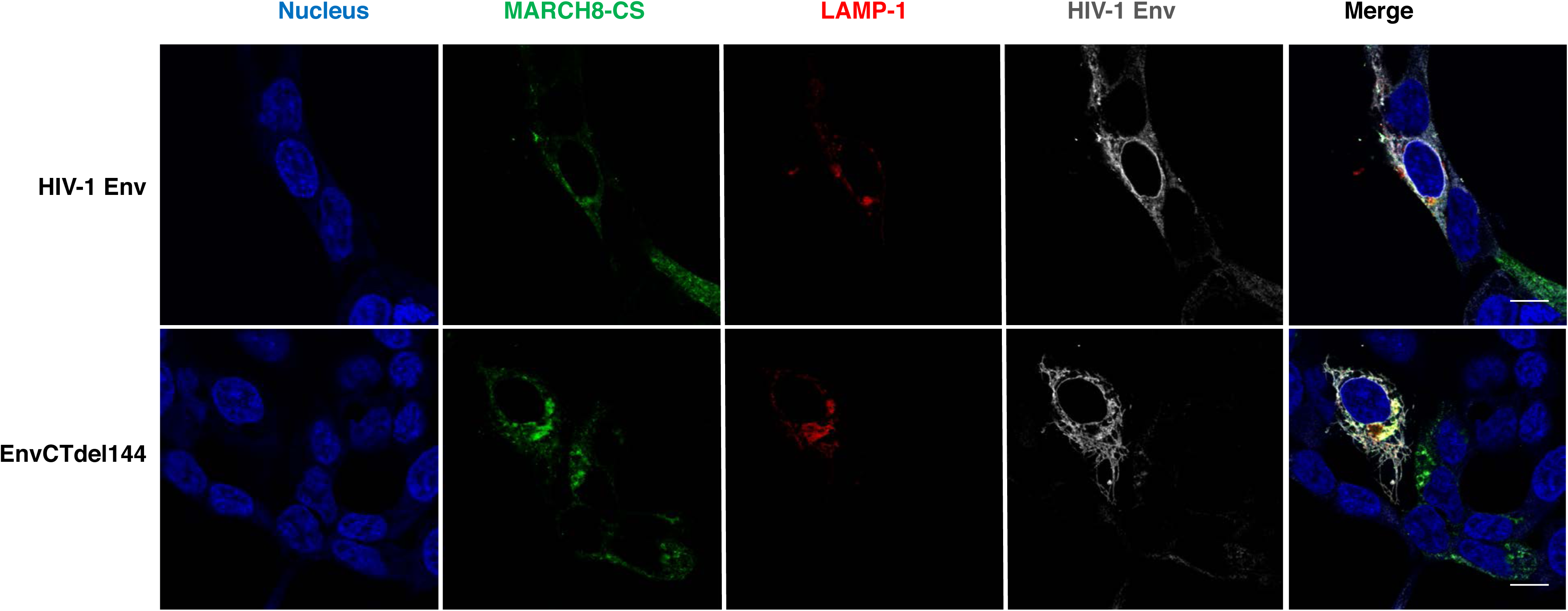
MARCH8 traps WT and truncated HIV-1 Env in an intracellular LAMP-1^+^ compartment. HEK293T cells were cotransfected with vectors expressing WT or truncated HIV-1 Env (pIIINL4-Env or pIIINL4-EnvCTdel144, respectively) (200 ng), pSV-Tat, and HA-tagged MARCH8 expression vectors (100 ng) and the LAMP1-RFP or pDsRed-Golgi-Beta 1,4- galactosyltransferase (300 ng) expression vectors. One day post-transfection, cells were processed for confocal microscopy. Distribution of WT HIV-1 Env **(A)**, and EnvCTdel144 **(B)** in the absence or presence of WT MARCH8 or MARCH8-CS. Colocalization of HIV-1 Env and EnvCTdel144 with MARCH8 **(C)** and MARCH-CS **(D)** and the LAMP1-RFP or pDsRed-Golgi-Beta 1,4- galactosyltransferase cellular markers. Scale bars = 10 µm.

The confocal microscopy data obtained with EboV-GP and SARS-CoV-2 S protein paralleled our observations with HIV-1 Env but diverged from the results obtained with VSV-G. Expression of MARCH8 but not MARCH8-CS reduced the expression of these viral glycoproteins on the cell-surface and induced their retention in an internal, LAMP-1^+^ compartment. The localization of truncated and K-to-R EboV-GP and S protein mutants was also shifted by MARCH8 expression (**Figs. 11 and 12; Table 1**).

**Figure 11.**
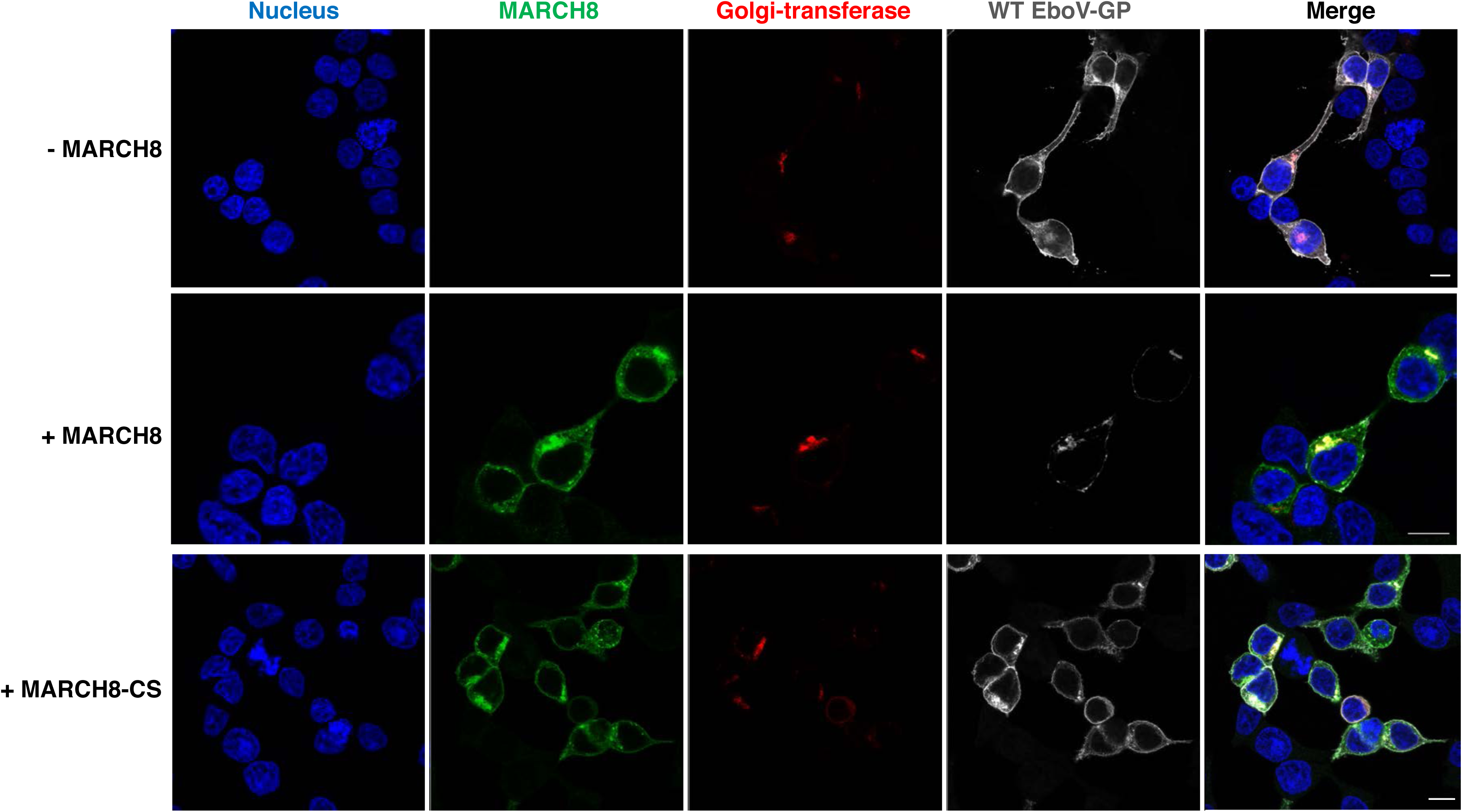
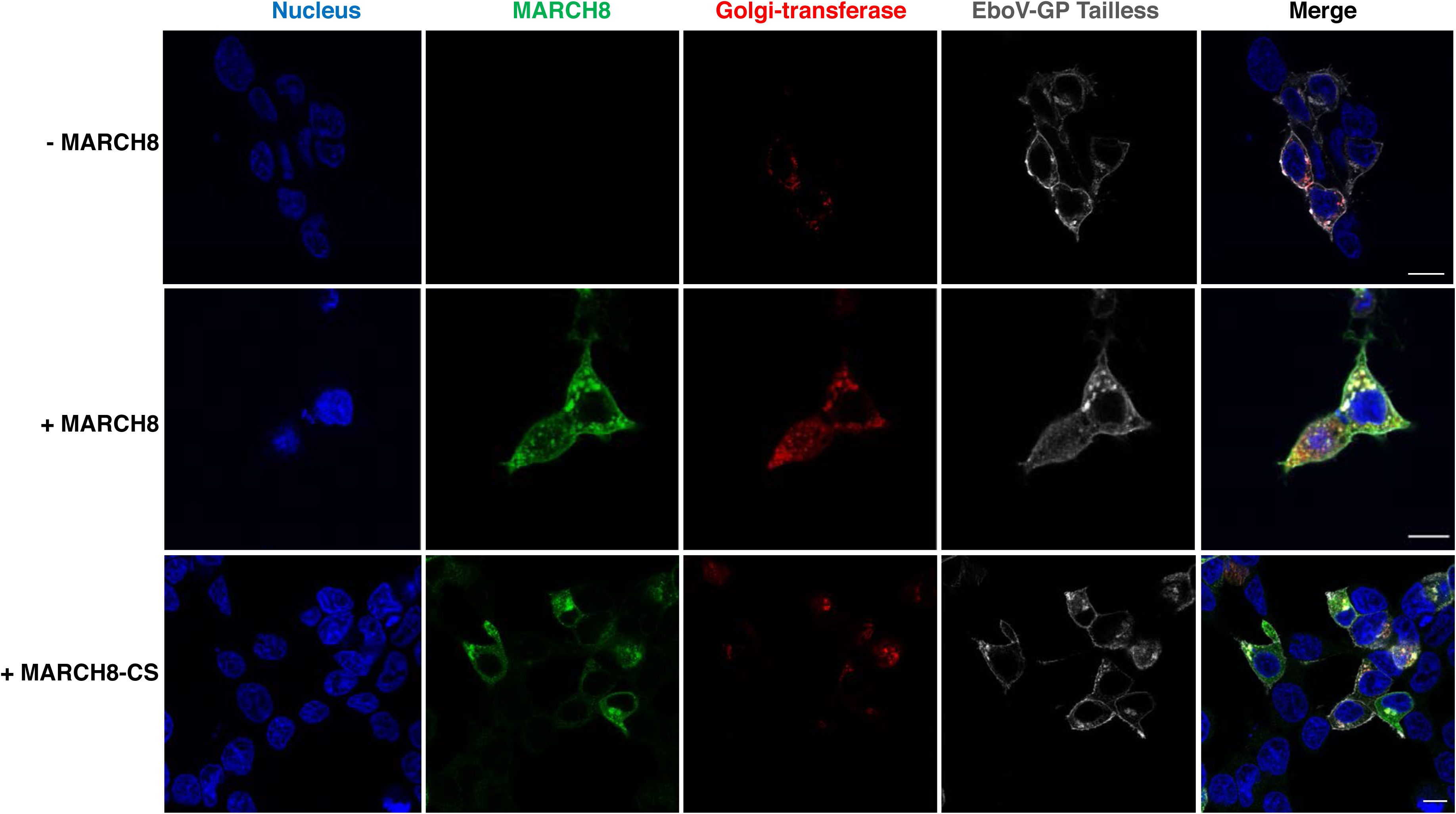
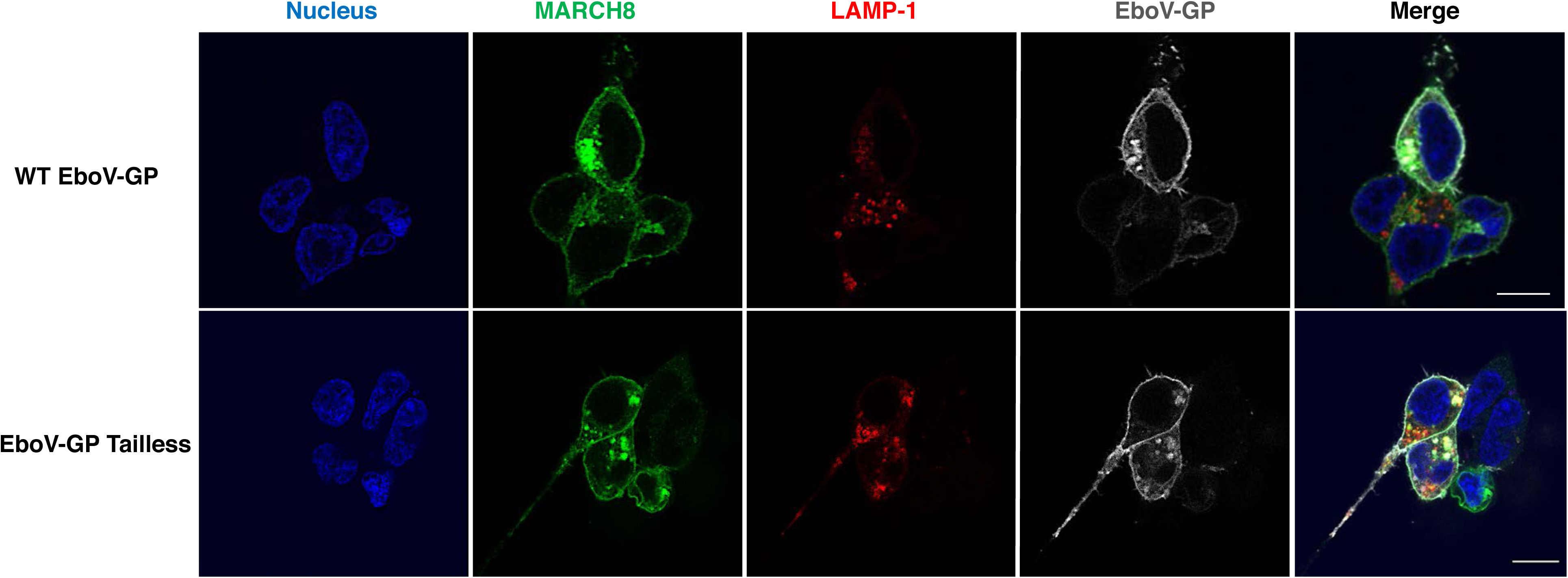
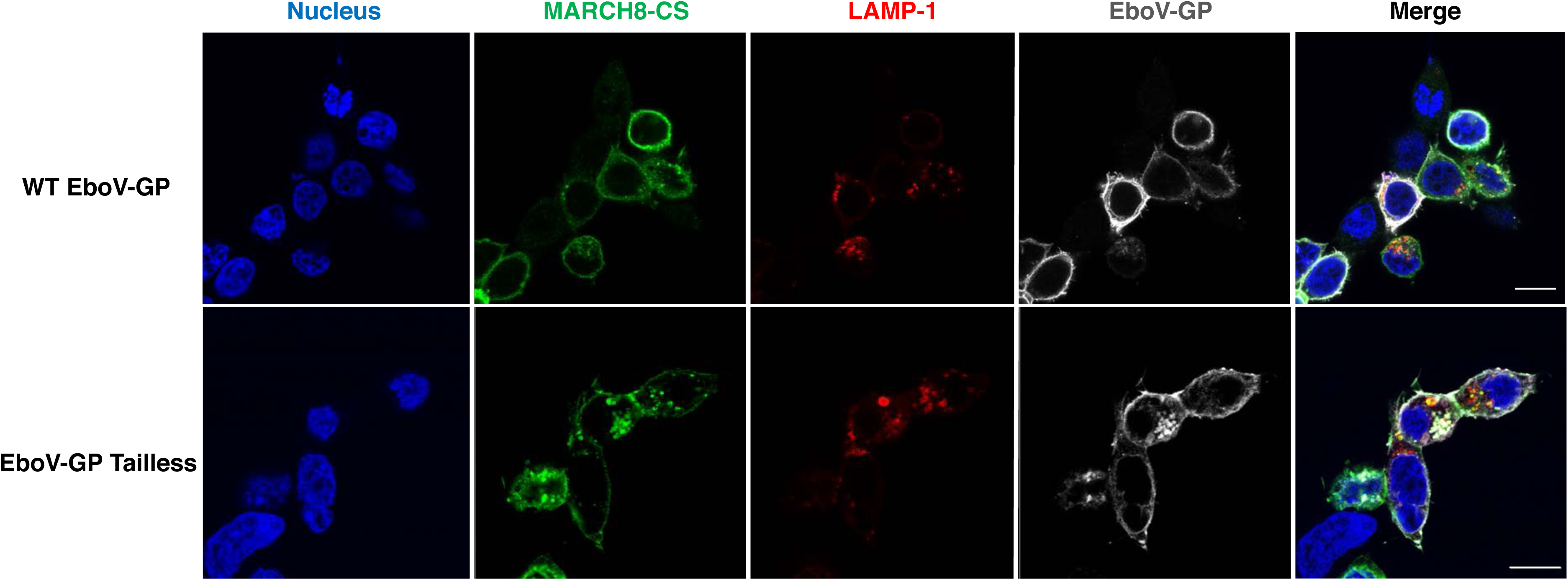
MARCH8 traps WT and truncated EboV-GP in an intracellular LAMP-1^+^ compartment. HEK293T cells were cotransfected with vectors expressing EboV-GP or EboV-GP Tailless (200 ng) and HA-tagged MARCH8 expression vectors (100 ng) and the LAMP1-RFP or pDsRed-Golgi-Beta 1,4-galactosyltransferase (300 ng) expression vectors. One day post-transfection, cells were processed for confocal microscopy. Distribution of EboV-GP **(A)** and EboV-GP Tailless **(B)** in the absence or presence of WT MARCH8 or MARCH8-CS. Colocalization of EboV-GP and EboV-GP Tailless with MARCH8 **(C)** and MARCH8-CS **(D)** and the LAMP1-RFP or pDsRed-Golgi-Beta 1,4-galactosyltransferase cellular markers. Scale bars = 10 µm.

**Figure 12.**
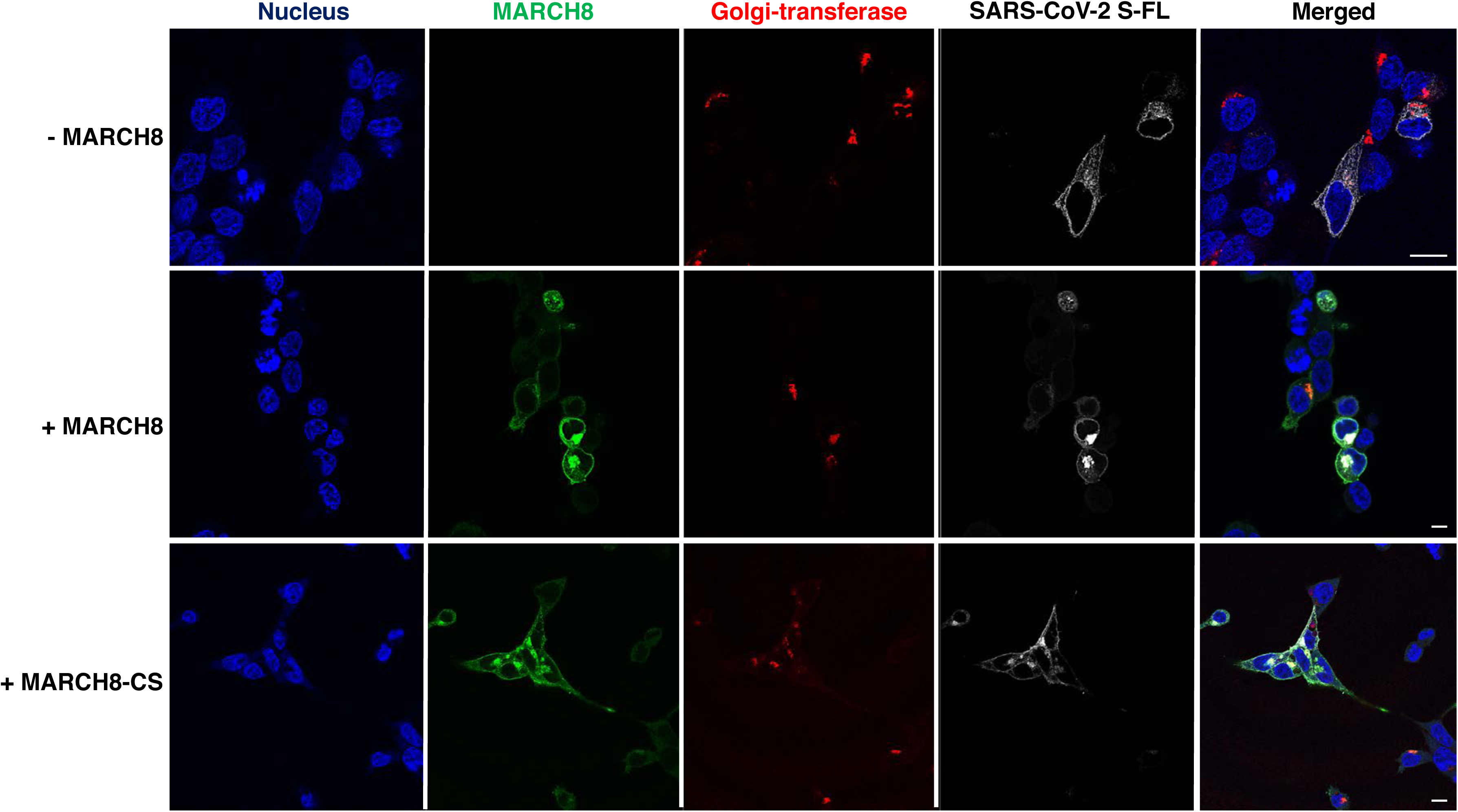
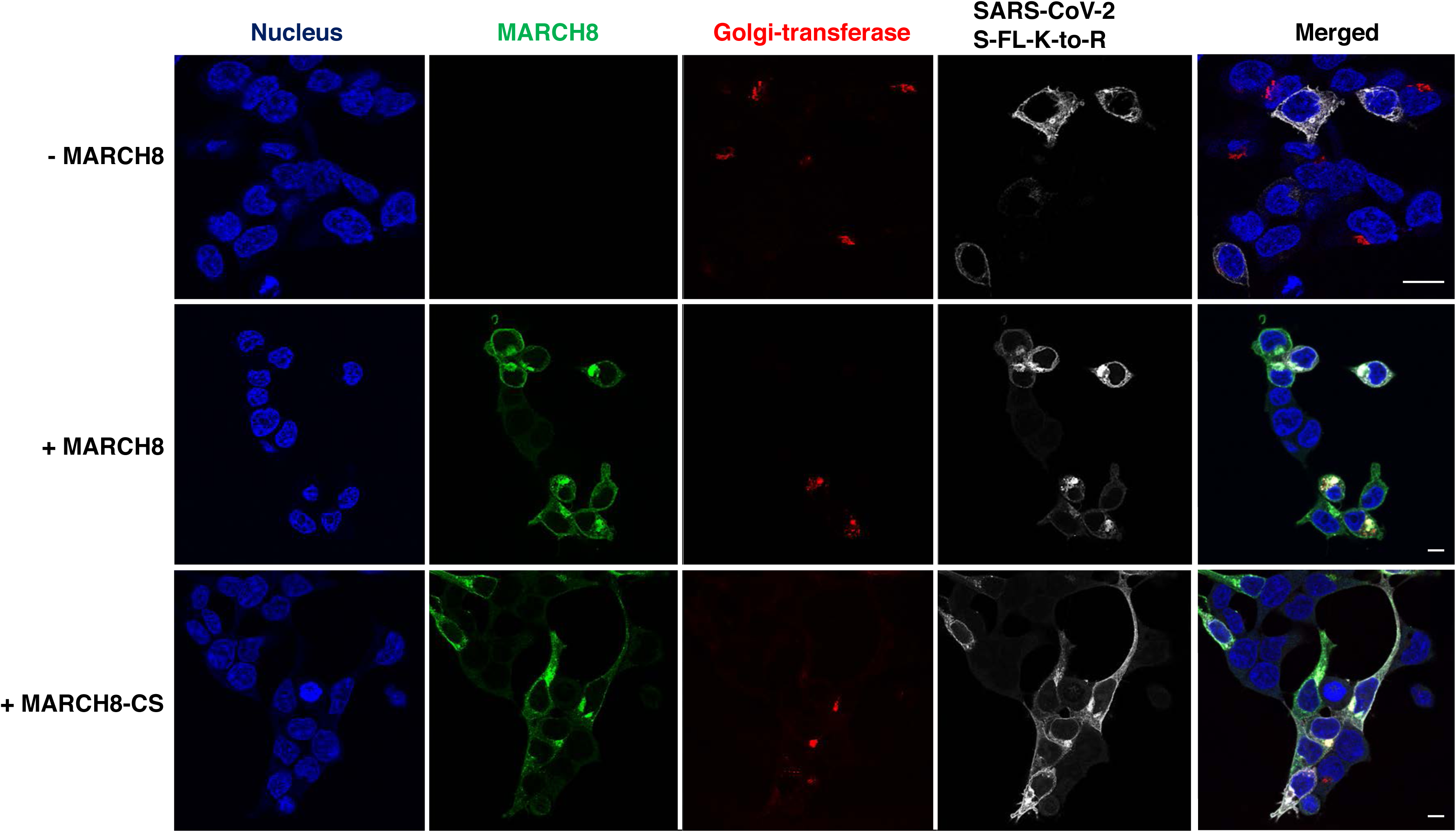
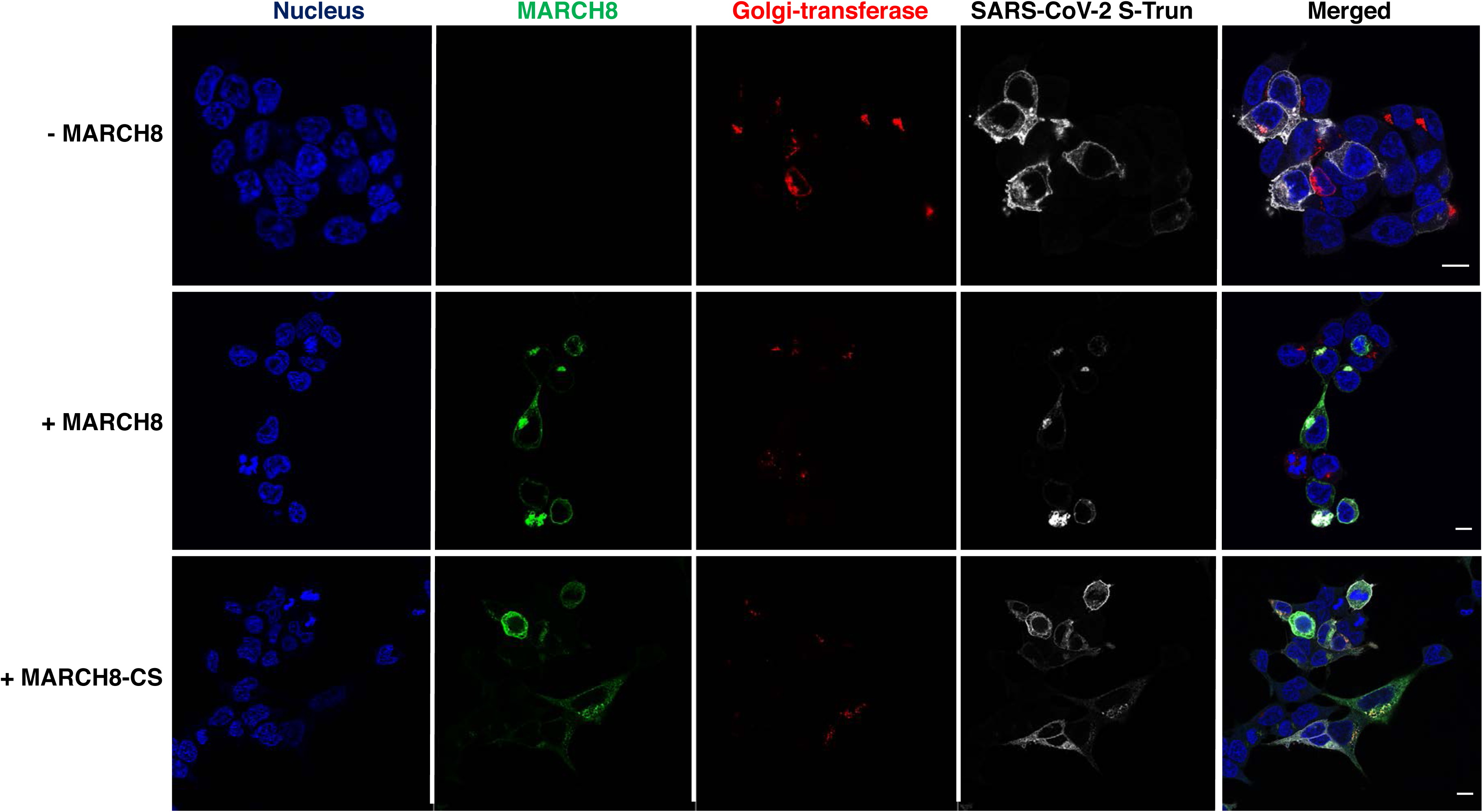
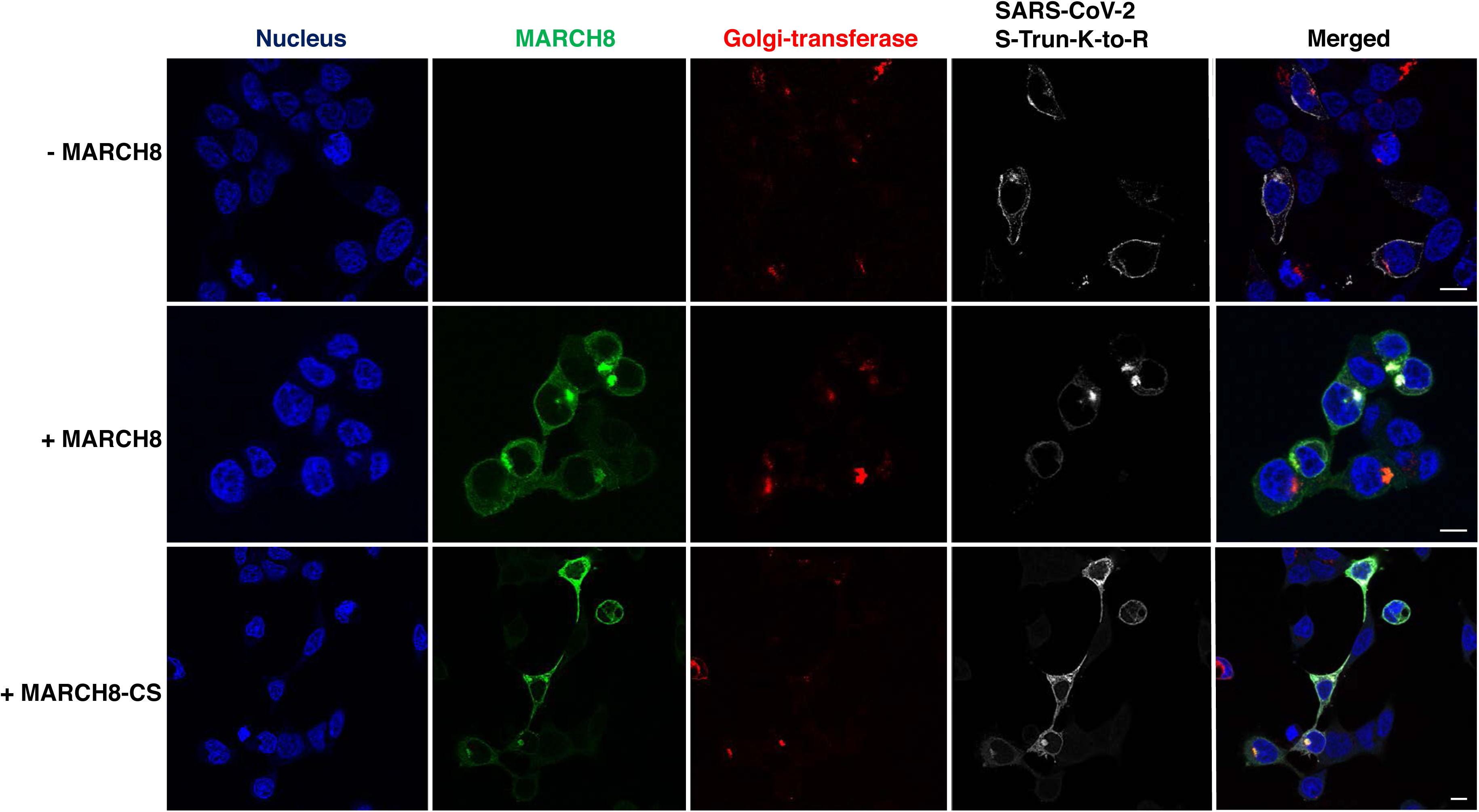
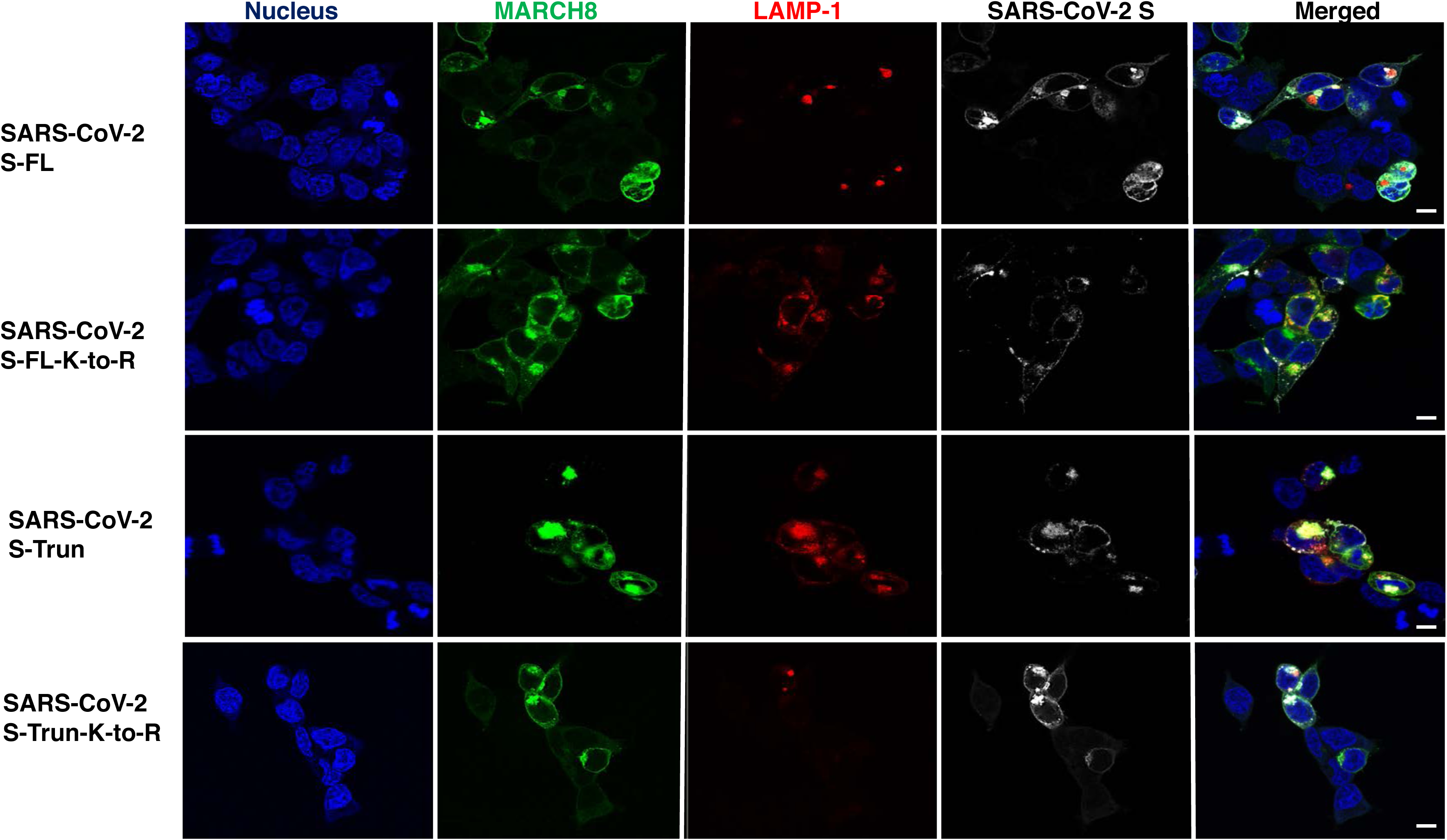
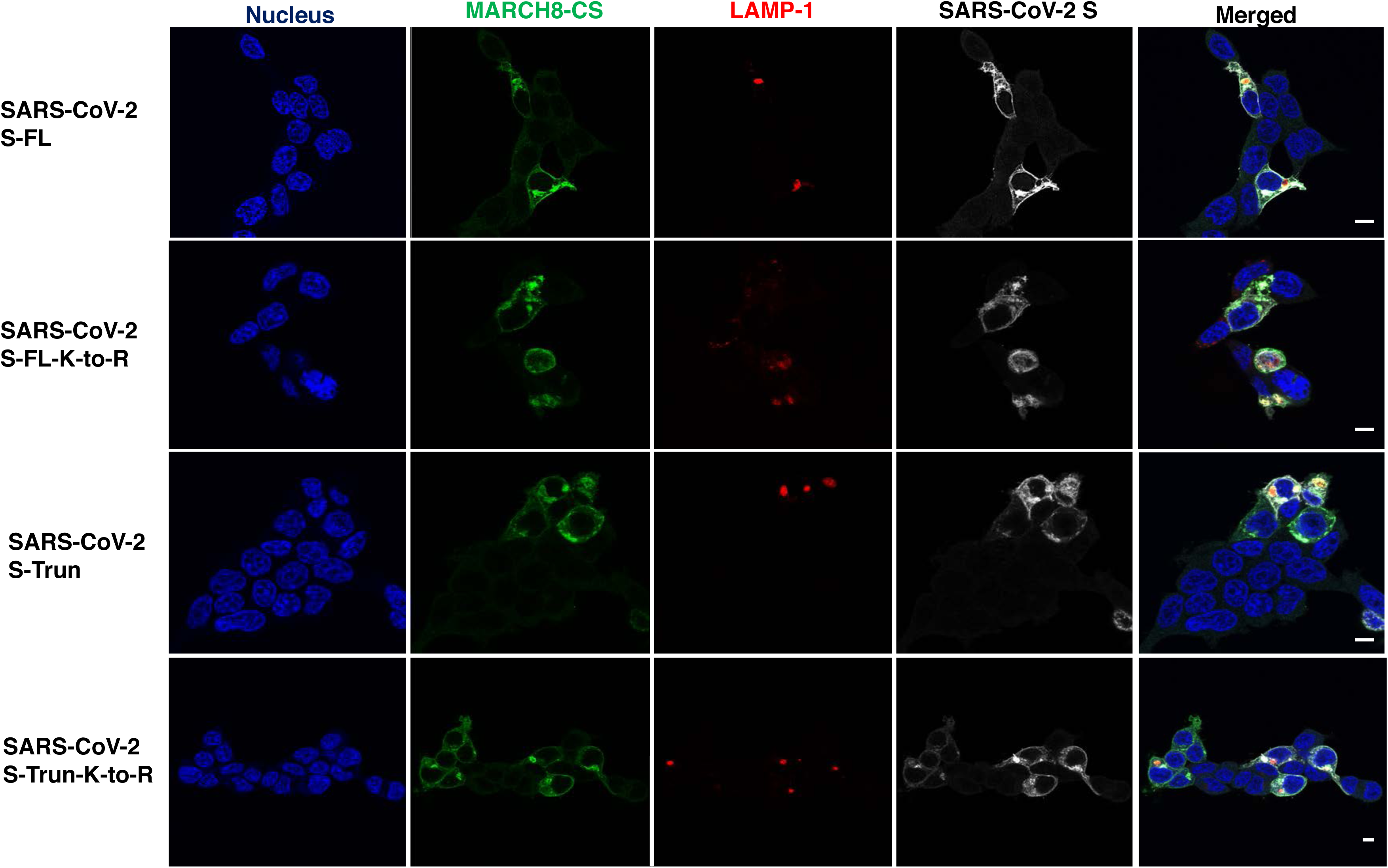
MARCH8 traps WT and CT-mutant SARS-CoV-2 S proteins in an intracellular LAMP-1^+^ compartment. HEK293T cells were cotransfected with vectors expressing WT or CT- mutant SARS-CoV-2 S proteins (200 ng) and HA-tagged MARCH8 expression vectors (100 ng) and the LAMP1-RFP or pDsRed-Golgi-Beta 1,4-galactosyltransferase (300 ng) expression vectors. One day post-transfection, cells were processed for confocal microscopy. Distribution of SARS-CoV-2 S-FL **(A)**, SARS-CoV-2 S-FL-K-to-R **(B)**, and SARS-CoV-2 S-Trun **(C)** and SARS-CoV-2 S-Trun-K-to-R **(D)** in the presence of MARCH8 and MARCH8-CS. Colocalization of SARS-CoV-2 S WT and CT mutants with MARCH8 **(E)** and MARCH8-CS **(F)** and the LAMP1-RFP or pDsRed-Golgi-Beta 1,4-galactosyltransferase cellular markers. Scale bars = 10 µm.

Taken together, the confocal microscopy data demonstrate that MARCH8 expression redirects the localization of viral glycoproteins by two distinct mechanisms. In the case of VSV-G, the CT is required for the MARCH8-induced relocalization to a LAMP-1^+^ compartment and mutation of the Lys residues in the CT abrogates the ability of MARCH8 expression to disrupt the localization of the viral glycoprotein. In contrast, for HIV-1 Env, EboV-GP and SARS-CoV-2 S protein, neither the CT nor the Lys residues in the CT are required for MARCH8-induced relocalization. The microscopy data are thus consistent with the infectivity and western blotting data (**Fig. 2-6**) showing that the ability of MARCH8 to antagonize VSV-G is dependent on the CT but the antagonism of the other viral glycoproteins tested is CT independent.

## DISCUSSION

In this study, we examined the ability of MARCH8 to antagonize a wide range of viral envelope glycoproteins – specifically, HIV-1 Env, VSV-G, EboV-GP and SARS-CoV-2 S protein. Expression of MARCH8 in the virus-producer cell reduced levels of viral glycoprotein processing for those glycoproteins that undergo furin-mediated cleavage, reduced glycoprotein stability, and interfered with glycoprotein incorporation into HIV-1 particles. In the case of VSV-G, MARCH8- mediated antagonism required the CT of the viral glycoprotein, and Lys residues in the CT were essential for MARCH8 restriction. In contrast, in the case of HIV-1 Env, EboV-GP, and SARS-CoV-2 S protein, the CT of the viral glycoprotein was not required for the antiviral effect. In all cases, however, the ubiquitin ligase activity of MARCH8 was required for restriction, as RING domain mutations to a large extent abrogated the inhibition. While this manuscript was in preparation, another group reported that, as we observed here, MARCH8-mediated restriction of VSV-G was CT-dependent but antagonism of HIV-1 Env was CT-independent, and they demonstrated MARCH8-mediated ubiquitination of VSV-G (55). These results support a model whereby the RING domain E3 ubiquitin ligase activity of MARCH8 either directly targets the CT of the viral glycoprotein – as has been shown for MARCH protein-mediated downregulation of a variety of cellular proteins (18) – or exerts its effect indirectly. In either case, the consequence of MARCH8 activity is the apparent redirection of the viral glycoproteins to a LAMP-1^+^ compartment. This is consistent with MARCH8 inducing lysosomal degradation of viral glycoproteins.

While MARCH8 expression antagonized each of the glycoproteins tested here, the magnitude of the effect varied across the different glycoproteins, in terms of both particle infectivity and glycoprotein incorporation into virions. Specifically, the inhibition of VSV-G and HIV-1 Env was quite severe, whereas more modest effects were observed with EboV-GP and SARS-CoV-2 S. The degree of inhibition did not correlate with the viral glycoprotein incorporation efficiency; for example, EboV-GP was very efficiently incorporated into HIV-1 particles, whereas SARS-CoV-2 S protein was inefficiently incorporated. Further study will be required to determine the basis for the variable activity of MARCH8 against the four glycoproteins examined in this study.

It remains unclear at what stage during their trafficking pathway viral glycoproteins are targeted by MARCH8. MARCH8 could act early during trafficking of the viral glycoprotein through the Golgi *en route* to the site of virus assembly, at the plasma membrane, or, in the case of glycoproteins like HIV-1 Env that undergo a recycling step, could act late following endocytic uptake from the plasma membrane. MARCH proteins have been reported to target a large repertoire of cellular proteins, including innate/adaptive immune receptors and intracellular adhesion molecules (18). Previous studies have shown that the overexpression of MARCH proteins can cause redistribution of syntaxin 4 and syntaxin 6 as well as some syntaxin-6- interacting soluble *N*-ethylmaleimide-sensitive factor attachment protein receptors (SNAREs), which are known to be involved in cellular endosomal trafficking (56, 57). MARCH protein overexpression has also been reported to alter the trafficking of clathrin-independent endocytosis (CIE) cargo proteins, rerouting them to endosomes and lysosomes for degradation (29). Thus, the indirect mechanism of MARCH8-mediated targeting of viral glycoproteins that we hypothesize would involve the ubiquitination and downregulation of cellular factor(s) involved in viral glycoprotein trafficking. The resulting accumulation of the viral glycoprotein in a LAMP-1^+^ intracellular compartment (Figs. 9-12), would reduce virion incorporation of the viral glycoprotein and impair particle infectivity. Taken together, our data suggest that MARCH8 exerts its antiviral activity via two different mechanisms, one (e.g., as observed for VSV-G) in which the CT of the viral glycoprotein is directly ubiquitinated, the other (e.g., as observed for HIV-1 Env, EboV-GP, and SARS-CoV-2 S) in which host trafficking factor(s) are targeted, resulting in an indirect restriction of the viral glycoprotein.

The viral homologs of the cellular MARCH proteins have been reported to ubiquitinate residues other than Lys in the CTs of their target proteins; for example, the K3 protein of KSHV has been reported to attach ubiquitin to Cys residues and the mK3 protein of gamma herpesviruses can ubiquitinate Ser and Thr in addition to Lys (35–37, 58). It is not clear whether MARCH proteins are able to ubiquitinate non-Lys residues, but we cannot exclude this possibility. Inspection of the viral glycoproteins under study indicates that the truncated HIV-1 Env mutant CTdel144 and EboV-GP Tailless have no Ser, Thr or Cys residues facing the cytosol, indicating that their CTs cannot be ubiquitinated. In contrast, SARS-CoV-2 S-Trun has 4 Ser, 1 Thr and 10 Cys residues. Ongoing work is investigating whether the CT of SARS-CoV-2 S protein undergoes MARCH8-mediated ubiquitination. It is interesting to note that the basic residues in the CT of HIV-1 gp41 are highly skewed towards Arg instead of Lys residues; for example, the CT of the NL4-3 strain of HIV-1 used in this study contains 21 Arg residues but only two Lys residues (https://www.hiv.lanl.gov/content/sequence/HIV/mainpage.html). This observation invites the speculation that HIV-1 may have evolved to limit the number of Lys residues available for gp41 CT ubiquitination.

Our results show a broad antiviral activity of MARCH8 against the glycoproteins from three human viral pathogens – HIV-1, EboV, and SARS-CoV-2 – and a primarily animal pathogen, VSV. MARCH8 targeting of HIV-1 Env, VSV-G, and, very recently, EboV-GP, has been reported (6, 8, 11, 55, 59). HIV-1 replicates predominantly in CD4^+^ T cells and also infects MDM (60). EboV targets mainly alveolar macrophages and endothelial cells (61); SARS-CoV-2, like other coronaviruses, targets airway epithelial cells (62); and VSV typically replicates in cells of the oral mucosa, although VSV-G confers very broad tissue tropism (63). In this study, we relied extensively on expressing MARCH8 exogenously to evaluate its effect on viral envelope glycoprotein stability, trafficking, and incorporation. This raises the question of whether the levels of MARCH8 expression achieved in these experiments are physiologically relevant. A previous study demonstrated that depletion of MARCH8 in primary human MDM significantly increased the infectivity of HIV-1 particles released from these cells, suggesting that at least in this cell type endogenous levels of MARCH8 expression are sufficient to restrict HIV-1 infectivity (11). MARCH8 expression is high in the lung (19), suggesting its potential relevance to infection by respiratory viruses like SARS-CoV-2. Due to the poor quality of available MARCH8 antibodies, we could not evaluate endogenous MARCH8 protein expression. Instead, qRT-PCR was used to measure the endogenous *MARCH8* gene expression in relevant cell types in the presence and absence of IFN. In the absence of IFN stimulation, basal levels of *MARCH8* RNA, measured as a ratio to *GAPDH*, were similar across the cell types analyzed. The RNA levels measured following IFN induction were comparable to those in transiently transfected HEK293T cells. Although RNA levels may not be directly correlated with protein expression, the results suggest that the MARCH8 expression in the cell types tested may be sufficient to exert an antiviral activity. Indeed, even in the absence of IFN stimulation, we observed that depletion of MARCH8 in HEK293T cells caused a modest but statistically significant increase in the infectivity of particle produced from those cells. In addition, our data indicate that *MARCH8* is an IFN-stimulated gene (ISG). The type I IFN (IFN-I) response is crucial in the host antiviral defense against viral infections (64, 65). A recent publication highlighted the importance of the IFN-I response in immune protection against SARS-COV-2 by establishing a link between life-threatening COVID-19 symptoms and loss-of-function mutations in IFN-I-related genes (66).

Viruses, including those whose envelope glycoproteins were studied here, have evolved a complex array of countermeasures to disable the function of cellular ISGs (67) . Further work will be required to elucidate mechanisms by which viruses counteract the antiviral activity of the MARCH family of E3 ubiquitin ligases.

## Materials and Methods

### Cell culture

HEK293T cells [obtained from the American Type Culture Collection (ATCC) and TZM-bl cells (obtained from J. C. Kappes, X. Wu, and Tranzyme, Inc. through the NIH AIDS Reagent Program (ARP), Germantown, MD)] were maintained in DMEM containing 5% or 10% (vol/vol) fetal bovine serum (FBS; HyClone), 2mM glutamine, 1% penicillin-streptomycin (penicillin 50 U/ml and streptomycin 50 μg/ml, final concentration; Lonza) at 37 °C with 5% CO_2_. The SupT1 T-cell line was cultured in RPMI with 10% FBS and 1% penicillin-streptomycin. The A549 human lung epithelial cell line (obtained from the ATCC) was cultured in DMEM with 10% FBS and 1% penicillin-streptomycin. Primary small airway epithelial cells (obtained from ATCC) were cultured in airway epithelial cell basal medium (ATCC) supplemented with bronchial epithelial cell growth kit (ATCC). Primary hPBMC were obtained from healthy volunteers from the NCI-Frederick Research Donor Program. The hPBMCs were extracted from whole blood using the Histopaque procedure (Sigma). Cells were then stimulated with PHA-P and IL-2 for three days.

### Plasmids and transfection

The following plasmids were used in this study: the full-length HIV-1 clade B molecular clone pNL4-3 and derivatives pNL4-3EnvCTdel144 (68), pNL4-3KFS (*env*-minus) (69), pNL4-3.Luc.R-E- (NIH ARP, #3418); and the pHCMV-G (VSV-G expression vector) (70, 71). The EboV-GP ΔMLD was a gift from Judith White (University of Virginia) (47) and SARS-CoV-2 S protein (FL and Truncated) expression vectors were gifts from Thomas Gallagher (Loyola University). The MARCH8 and MARCH8-CS plasmids were gifts from Kenzo Tokunaga (National Institute of Infectious Diseases, Tokyo). Viral glycoprotein mutant plasmids VSV-G Tailless, VSV-G K-to-A and VSV-G K-to-R; EboV-GP K-to-A and EboV-GP-Tailless; SARS-CoV-2 S-FL K-to-R and SARS-CoV-2 S-Truncated K-to-R) were generated using the Q5® site-directed mutagenesis kit (New England BioLabs Inc., #E0552S) or the QuickChangeSite-Directed Mutagenesis kit (Stratagene) according to the manufacturer’s instructions with primers listed in Table S1. In microscopy experiments, pDsRed-monomer-Golgi-Beta-1,4- galactosyltransferase (Clontech, #632480) and LAMP-RFP (Addgene, #1817) (72) were used. Plasmids were purified using MaxiPrep Kits (Qiagen, #12263) and mutations were verified by sequencing (Psomagen, Rockville, MD). The HEK293T cells were transfected with the indicated plasmids using PEI or Lipofectamine 2000 (Invitrogen) according to manufacturer’s instructions.

Virus-containing supernatants were filtered through 0.45-μm membrane 24 or 48 hours post-transfection and virus was quantified by measuring RT activity. Virus-containing supernatant was harvested 24 or 48 hours post-transfection and virus particles were collected by ultracentrifugation. Virus and cell pellets were solubilized in lysis buffer [10mM iodoacetamide (Sigma-Aldrich), Complete^TM^ protease inhibitor tablets (Roche), 300 mM sodium chloride, 50mM Tris-HCl (pH 7.5) and 0.5% Triton X-100 (Sigma-Aldrich)] and used for further analysis.

### Western blotting

Cell and virus lysates in lysis buffer with 6x SDS-PAGE sample loading buffer (600 mM Tris-HCL pH6.8, 30% glycerol, 12% SDS, 20mM DTT, 0.03% bromophenol blue) were heated at 95°C for 5 min. Samples were analyzed on 10% 1.5mm Tris-glycine gels using a BioRad Trans-Blot Turbo Transfer system according to manufacturer’s instructions. Proteins were detected with primary (see Table S2) and secondary antibodies. Protein bands were visualized using chemiluminescence with either a Gel Doc XR+ system (BioRad) or Sapphire Biomolecular Imager (Azure Biosystems) and analyzed with either Image Lab version 6.0.1 or AzureSpot (Azure Biosystems).

### Single-cycle infectivity assays

TZM-bl is a HeLa-derived cell line that contains a stably integrated HIV-LTR-luciferase construct (73, 74). TZM-bl cells were infected with serial dilutions of RT-normalized virus stock (44) in the presence of 10 μg/ml DEAE-dextran. For SARS-CoV-2 S infectivity assays, HEK293T cells stably expressing hACE2 (BEI resources, NR-52511) were infected with serial dilutions of RT-normalized S-pseudotyped, luciferase-expressing HIV-1 (pNL4-3.Luc.R-E-) virus stock in the presence of 10 μg/ml DEAE-dextran. Cells were lysed with BriteLite Luciferase reagent (Perkin-Elmer) and luciferase was measured in a Wallac BetaMax Plate reader at 48 hours post-infection. Data were normalized to non-MARCH transfected negative control from three independent experiments.

### Confocal microscopy

Microscopy experiments used 18mm coverslips (Electron Microscopy Sciences, catalog number 72291-06) which were sterilized by soaking in ethanol and air-dried before seeding of HEK293T cells. Coverslips were pre-treated with fibronectin (Sigma-Aldrich, #GC010) at 1:50 dilution in PBS for 30 min at room temperature before seeding cells. HEK293T cells were cultured overnight and fixed with 4% paraformaldehyde (Electron Microscopy Sciences, BM-155) at 24 hr post-transfection in DPBS for 1 hr, quenched with 0.1M glycine in PBS for 10 min. Cells were permeabilized with 0.2% Triton X-100 (Sigma-Aldrich, #T-8787) in PBS for 5 min, blocked with 10% BSA/PBS (Sigma-Aldrich) for 30 min and stained with primary antibodies at a 1:400 dilution with 0.1% Triton and 1% BSA in PBS for 1 hr. After 3 washes in PBS, cells were incubated with secondary antibodies and DAPI stain in 1:1000 dilution in PBS. Antibody information is listed in Table S2. Cells were washed and mounted with Fluoromount-G (Electron Microscopy Sciences, #0100-01). Imaging was performed with a Leica TCS SP8 microscope (Leica Microsystems Inc., Buffalo Grove, IL) using a 63X oil-immersion objective. Images were generated using ImageJ software (NIH, Bethesda, MD). Background was subtracted using imageJ’s built-in ‘rolling ball’ background subtraction process. Colocalization analyses were performed with intracellular region of interest (ROI) excluding any plasma-membrane-associated signal using Colocalization test and a Plugin, EzColocalization, (75) with ImageJ. Untransfected cells were excluded from analysis.

### RT-qPCR

To measure *MARCH8* expression levels, total RNA was extracted using RNeasy Plus Mini Kit (Qiagen) following manufacturer’s instructions from transfected HEK293T, cell-lines or cells stimulated with 1000U/ml of IFN-α, β or γ (Pblassay science, catalog number 11200, 11415 and 11500, respectively). RNA was transcribed using High-Capacity cDNA Reverse Transcription Kit (Applied Biosystems) and real time PCR was performed using KAPA Sybr Fast master mix (Kapa Biosystems) following manufacturer’s instructions using CFX connect real-time PCR detection system (BioRad) with specific oligonucleotides described previously (6, 11). The *MARCH8* mRNA levels were normalized with *GAPDH* mRNA levels using ΔΔCT method for relative quantification.

### siRNA knockdown of endogenous *MARCH8*

HEK293T cells were transfected with 0.01, 0.1 and 1 μM MARCH8-specific siRNA or non-targeting control (NTC) (Smart pool, Dharmacon) using Lipofectamine RNAiMAX (ThermoFisher Scientific) following the manufactuer’s protocol. Seventy-two hrs post siRNA treatment, cells were collected for RT-qPCR to measure *MARCH8* expression. The siRNA-treated HEK293T cells were transfected with the HIV-1 pNL4-3 clone, and the progeny viruses were collected at 48 hr post-transfection, RT normalized and used to infect the TZM-bl indicator cell line.

## Acknowledgements

We thank Kenzo Tokunaga, Judith White, and Thomas Gallagher for providing plasmids for the study. We thank Sherimay Ablan and Melissa V. Fernandez for technical advice and members of the Freed lab for critical review of the manuscript and helpful discussion. We thank Kim Peifley and David Scheiblin (NCI-Frederick) for technical advice on microscopy.

## Author contributions

Designed research: CML, AW & EOF

Performed research: CML, AW, AM and NP

Analyzed data: CML & AW

Wrote paper: CML, AW and EOF

## Figure legends

**Figure S1. MARCH8 is localized in the endolysosomal compartments and at the plasma membrane.** HEK293T cells were cotransfected with MARCH8 expression vector (200 ng) and the LAMP1-RFP or pDsRed-Golgi-Beta 1,4-galactosyltransferase (300 ng) expression vectors. Cells were fixed, processed and stained with anti-HA to detect MARCH8. Scale bars = 10 µm. Images were acquired with a Leica TCS SP8 confocal microscope. Colocalization between MARCH8 and subcellular compartment markers was assessed by calculating the Pearson’s correlation coefficient (r values) ± SD using 50 cells per condition.

**Fig S2. Incorporation efficiency of viral glycoproteins into HIV-1 particles.** HEK293T cells were transfected with the WT (pNL4-3) or cotransfected with the Env-defective (pNL4-3/KFS) HIV-1 molecular clone and vectors expressing VSV-G or EboV-GP proteins or the Env-defective, luciferase-expressing pNL4-3 derivative pNL4-3.Luc.R-E- and a vector expressing the SARS-CoV-2 S protein. Two days post-transfection, cell and viral lysates were prepared and subjected to western blot analysis with antibodies against gp41, VSV-G, EboV GP2, or SARS-CoV-2 S2. The levels of viral glycoprotein in cell and virus lysates were quantified and incorporation efficiency was calculated as the amount of virion-associated glycoprotein relative to total glycoprotein in cell and virus. Data shown are + SD from three independent experiments. Plasmid concentrations indicated in legends of Fig 3-6.

