## Supplemental Figs and Table S1 for "Mechanism of Viral Glycoprotein Targeting by Membrane-associated-RING-CH Proteins"

Fig. S1.

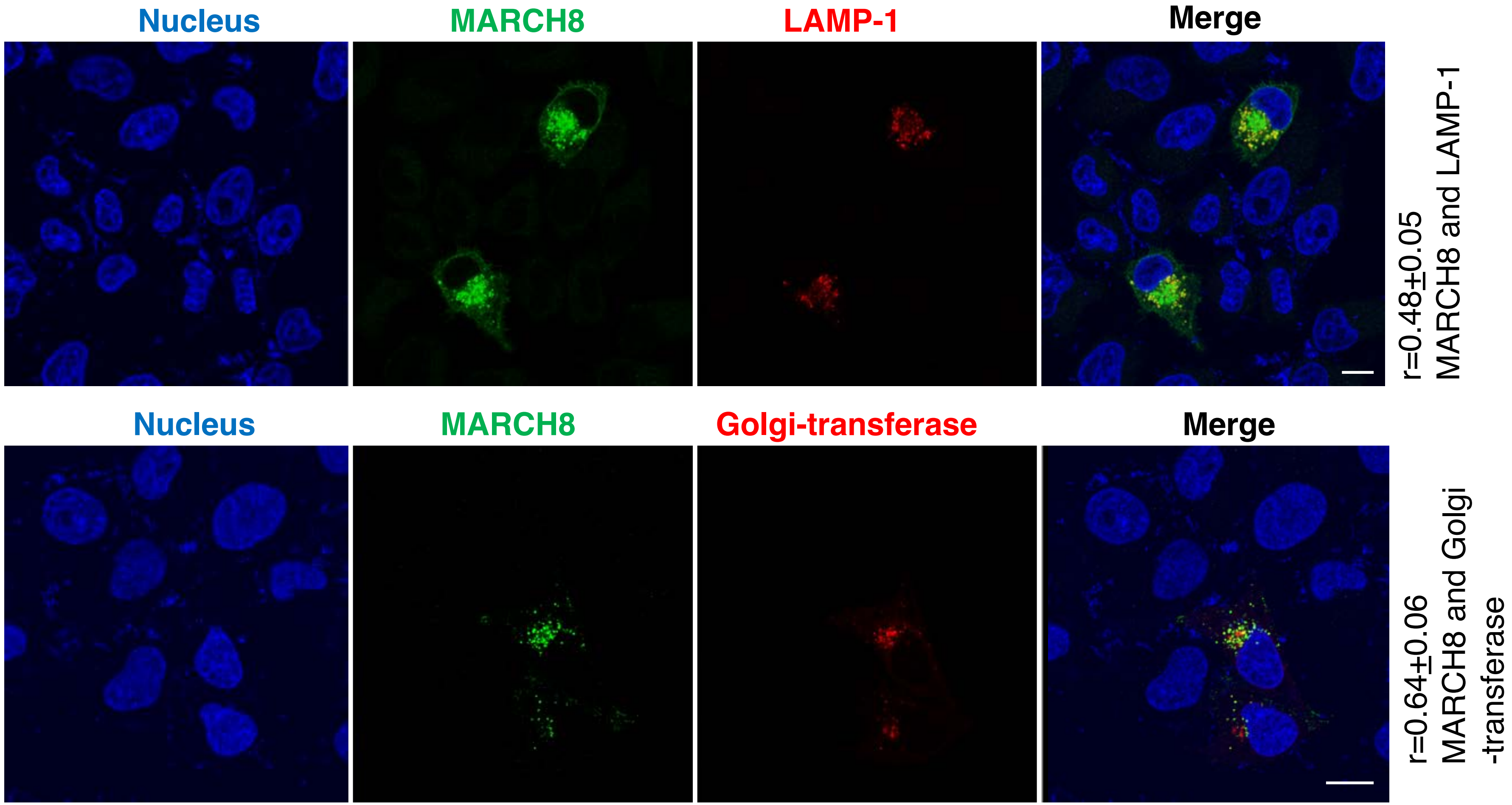
Nucleus
MARCH8
LAMP-1
Merge
r=0.48±0.05
MARCH8 and LAMP-1
Nucleus
MARCH8
Golgi-transferase
Merge
r=0.64±0.06
MARCH8 and Golgi
-transferase

Fig. S2.

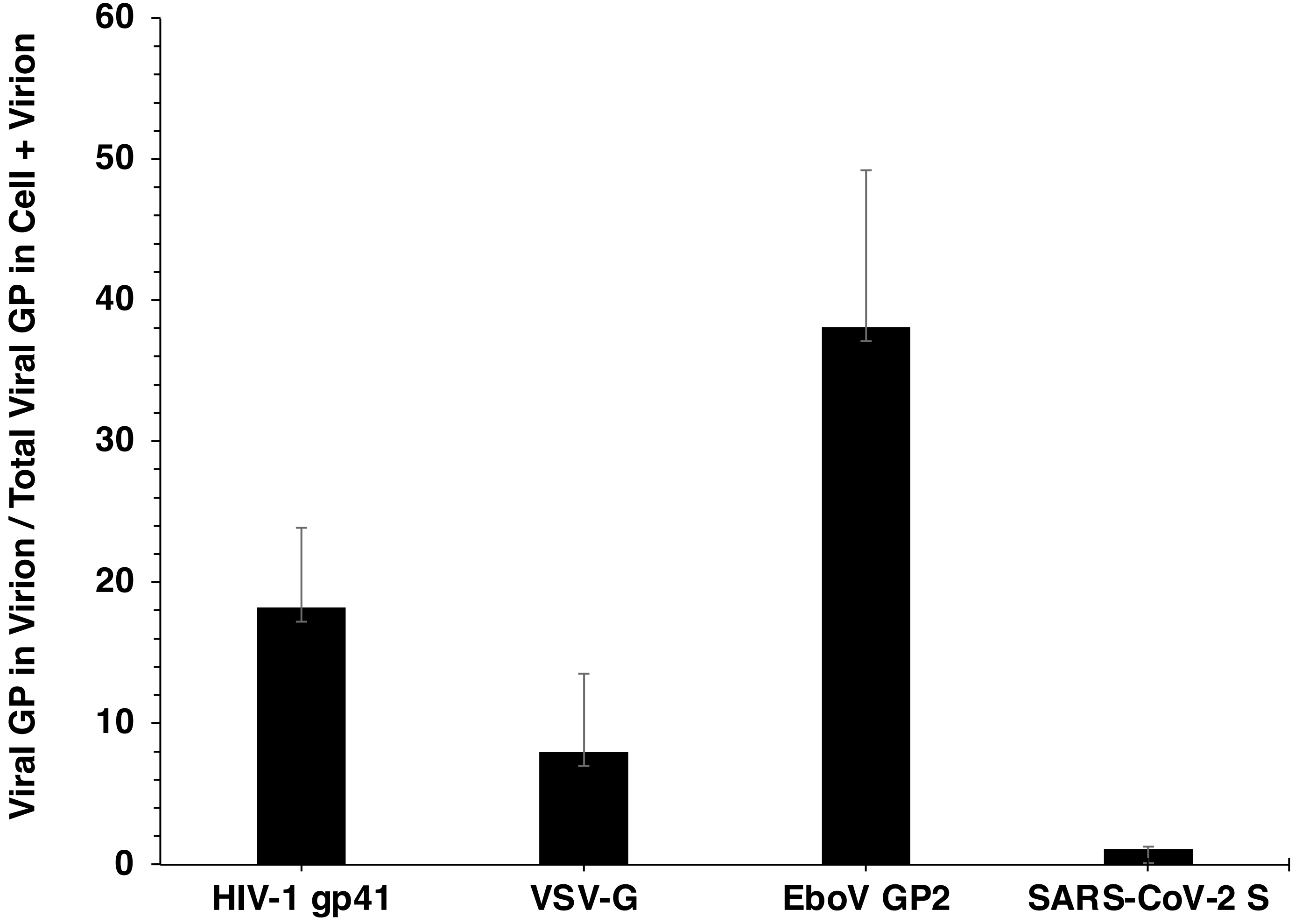
Viral GP in Virion / Total Viral GP in Cell + Virion
60
50
40
30
20
10
0
HIV-1 gp41
VSV-G
EboV GP2
SARS-CoV-2 S

Table S1.

Table S1. Basal expression of MARCH8 relative to GAPDH

| | GAPDH (Ct) [c] | MARCH8 (Ct) [c] | MARCH8/ GAPDH ratio |
|---|---|---|---|
| hPAECs [a] | 14.22 | 22.65 | 1.59 |
| SupT1 T cell line | 14.25 | 22.10 | 1.55 |
| hPBMCs [b]-Donor1 | 14.93 | 21.81 | 1.46 |
| hPBMCs-Donor2 | 14.88 | 22.94 | 1.54 |
| hPBMCs-Donor3 | 14.58 | 21.8 | 1.50 |
| HEK293T cell line | 14.23 | 22.04 | 1.55 |

[a] hPAECs, human primary airway epithelial cells

[b] hPMBCs, human peripheral blood mononuclear cells

[c] Cycle threshold (Ct) values for GAPDH and MARCH8. Data shown are Ct averages from three independent experiments for hPAECs, SupT1, and HEK293T
